# Multi-Omics Mapping of Human Kidney Reveals Complement-Mediated Cellular Dynamics During Progression of Focal Segmental Glomerulosclerosis

**DOI:** 10.64898/2026.02.19.706727

**Authors:** Sahomi Hayashi, Miyoshi Takeuchi, Toshiaki Nakano, Daiki Setoyama, Sasha A. Singh, Abhijeet R. Sonawane, Takaki Iwamoto, Hiroshi Kishimoto, Akihiro Tsuchimoto, Shunsuke Yamada, Dongchon Kang, Tetsuro Ago, Takanari Kitazono, Masanori Aikawa, Yuya Kunisaki

## Abstract

Focal segmental glomerulosclerosis (FSGS) is a major cause of glucocorticoid-resistant nephrosis, yet its pathogenesis remains unclear. To define the molecular and cellular landscape of FSGS, we employed multi-omics approaches on independent human kidney biopsy cohorts. Proteomics revealed enhanced immune and complement activation. Spatial transcriptomics using a target gene panel constructed from the proteomics highlighted alternative pathway activation driven by complement factor D as a prominent feature. Complement activation emerged in glomeruli alongside altered signaling in podocytes and parietal epithelial cells (PECs): Podocytes displayed reduced PECAM1 and elevated macrophage migration inhibitory factor (MIF) signaling, and PECs display complement and collagen signaling as early events, and progressively acquired cellular activation, prompting glomerular fibrosis. These pathogenic signals propagated into adjacent interstitial regions, paralleling immune cell infiltration and fibrosis. These results uncover dynamics of dysregulated intercellular interactions originated from podocyte and PECs underlying the pathogenesis of FSGS, which provides insights into potential stage-specific interventions.

## INTRODUCTION

Focal segmental glomerulosclerosis (FSGS) is a clinicopathological syndrome defined as extracellular matrix accumulation in some glomeruli (focal) and obstruction of glomerular capillaries affecting a portion of the glomerular tuft (segmental)^1^. It presents with proteinuria resulting from injury of podocytes, which are specialized epithelial cells responsible for forming glomerular filtration barriers^2^. Clinically, 40–70% of FSGS patients are resistant to glucocorticoids^3^ and proportion of progression to end-stage kidney disease is 2.3%-40%^4,5^. Based on its etiology and clinical features, FSGS is classified into primary, secondary, genetic, and undetermined subtypes^1,6^. Accumulating evidence suggests that immune disorders contribute to the underlying pathology of FSGS, as evidenced by reports that nephrotic syndrome is remitted after measles infection^7^, and that glucocorticoids, immunosuppressive agents, and B cell-targeting rituximab are effective^8^. The precise pathogenic mechanisms, however, have yet to be defined, which has driven our current research efforts.

Omics-based approaches are becoming increasingly prevalent in glomerular disease research, providing powerful tools to elucidate previously unidentified disease mechanisms. In FSGS, proteomics studies have implicated pathways related to podocyte stress responses^9^ and glucocorticoid resistance^10^. These findings underscore the potential of proteomics to advance our understanding of FSGS pathogenesis. However, most previous analyses have relied on whole kidney tissues or glomeruli obtained by laser microdissection, which inevitably mask cell type–specific molecular alterations. Although single-cell RNA sequencing (scRNA-seq) enables high-resolution transcriptomic profiling, it lacks spatial information, preventing investigation of focal and segmental lesions characteristic of FSGS. To overcome these limitations, spatial transcriptomics has developed, which allows us to visualize molecular changes on preserved histological architecture. Several studies have applied spatial transcriptomics to glomerular diseases including FSGS and demonstrated transcriptional signatures such as downregulated podocyte-associated genes^11,12,13^. However, disease-specific molecular pathways in FSGS have not sufficiently explored.

Harnessing the strengths of proteomics approaches, we first performed comprehensive proteomics profiling of human kidney biopsy specimens to identify candidate molecular pathways associated with FSGS pathogenesis. Guided by these findings, we then employed high-resolution spatial transcriptomics using the Xenium in Situ platform to validate and localize intercellular communications and key pathways on tissue architecture.

## Results

### Patient demographics and study design

This study analyzed two independent cohorts comprising a total of 94 participants using multi-omics approaches. The proteomics cohort consisted of 73 individuals, including 23 patients with FSGS, classified as primary (n = 9), secondary (n = 8), and undetermined cause (n = 6), healthy kidney transplantation donors (n = 14), and disease controls with minimal change disease (MCD) (n = 15) or IgA nephropathy (IgAN) (n = 21). The spatial transcriptomics cohort comprised 21 individuals, including primary or undetermined cause of FSGS (n = 15) and kidney transplantation donors (n = 6). Patients with a history of nephrotic syndrome who had received glucocorticoid treatment were defined as primary FSGS, those with non-nephrotic syndrome and a history of secondary FSGS such as obesity or a long history of hypertension were defined as secondary FSGS, and those with non-nephrotic syndrome and no history of secondary causes were defined as FSGS of undetermined cause. Clinical characteristics, treatments and electron microscopy findings for each cohort are summarized in Supplementary Tables 1–5.

### Proteomics analyses identify immune and complement pathways enriched in the FSGS group

We first conducted proteomics analyses to investigate molecular characteristics in kidney of FSGS. To ensure quality of the kidney biopsy specimen used for this analysis, we examined glomerular occupancy in the tissues. To this end, the serial tissue sections prepared for Liquid Chromatograph-Tandem Mass Spectrometry (LC-MS/MS) analysis underwent hematoxylin and eosin (HE) staining, revealing similar glomerular proportions across the four groups (Extended Data Fig. 1a).

Our proteomics platform identified 5,749 proteins across all groups, of which 4,200 proteins with two or more unique peptides were classified as high confidence (Supplementary Table 6). After excluding potential keratin contaminants, 4,168 proteins remained for downstream analyses, and their intensities were normalized across samples, as illustrated in Extended Data Fig. 1b. The identified proteins included several markers for podocytes such as CD2 associated protein (CD2AP) and synaptopodin (SYNPO), which are important in maintenance of podocyte functions (Fig. 1a).

**Fig. 1.**
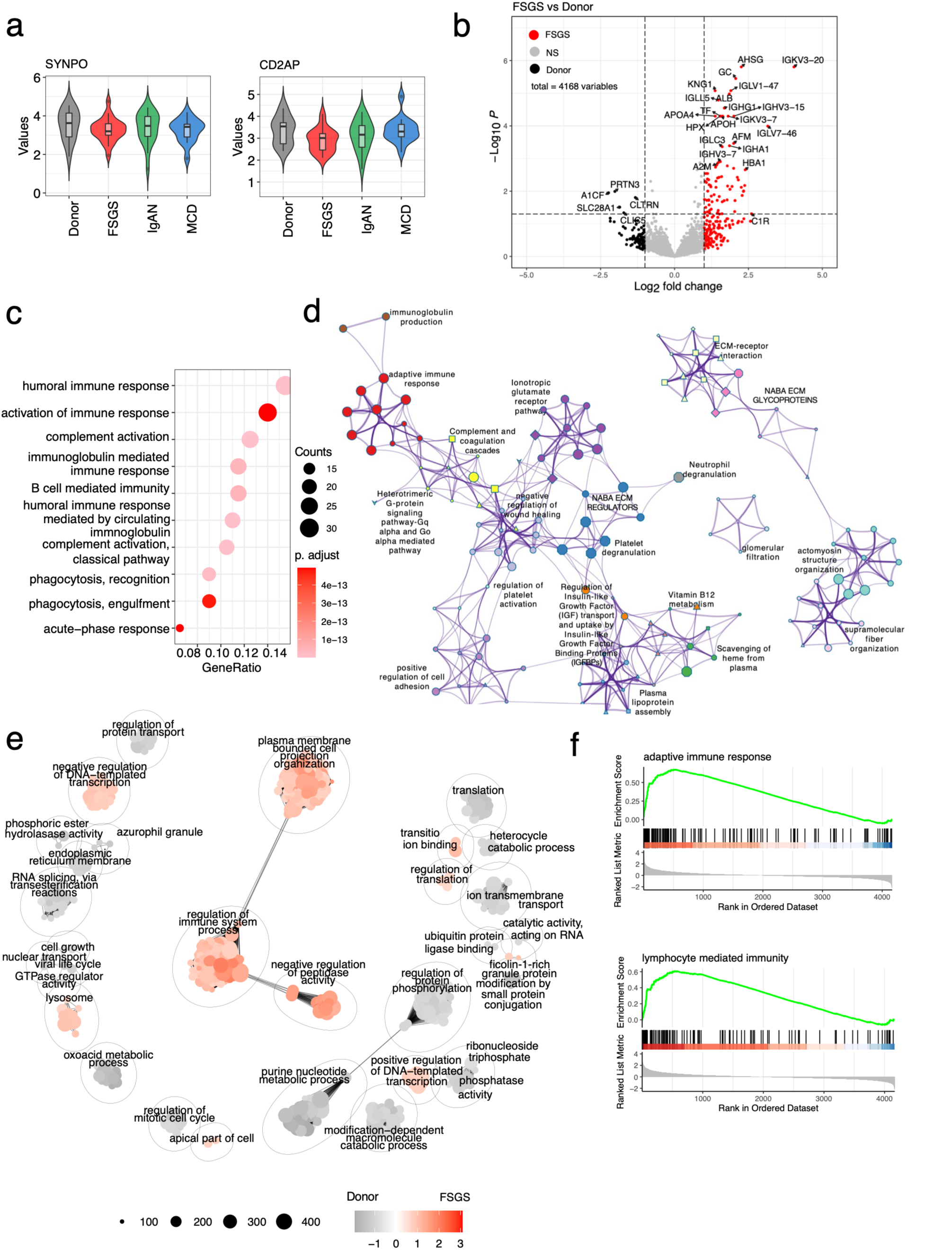
Proteomic profiling reveals enhanced immune responses and complement activation in FSGS. **a**, Violinplots of representative podocyte proteins, CD2 associated protein (CD2AP) and synaptopodin (SYNPO). **b,** A volcano plot of proteins abundant or reduced in the FSGS group. Significance was defined as P < 0.05 and |log2 fold change| > 1. NS, not significant **c,** Dot plot of Gene Ontology analysis of proteins increased in the FSGS group. Circle size indicates the number of annotated genes, and the red colour gradient indicates significance. **d,** Pathway enrichment analysis for the FSGS group compared with the kidney donor group using multiple databases. Shapes indicate pathway databases: Canonical pathways (diamond), Gene Ontology biological process (ellipse), KEGG (rectangle), Panther (V-shaped), Reactome (octagon), and WikiPathways (Triangle). **e,** GSEA for the FSGS group compared with the kidney donor group. Pathways enriched in the FSGS group are shown in red and those in the kidney donor group are shown in gray. Similar pathways are clustered together, with the circle size indicating the number of similar pathways. **f**, GSEA for adaptive immune response and lymphocyte mediated immunity in FSGS versus kidney donor.

Comparison between the FSGS and kidney donor groups identified 248 differentially abundant proteins (DAPs), with 175 proteins elevated and 73 reduced in the FSGS group (Fig. 1b, Supplementary Table 7a). Gene Ontology (GO) enrichment analysis of the upregulated DAPs in the FSGS group represented immune response, complement activation, immunoglobulin and B cell–mediated immunity and classical pathway (Fig. 1c, Extended Data Fig. 1c, Supplementary Table 7b). To examine functional relationships of the pathways enriched in the FSGS group, we conducted pathway network analysis using multiple databases including Canonical pathways, GO, Kyoto Encyclopedia of Genes and Genomes (KEGG), Panther, Reactome, and WikiPathways (Fig. 1d). The analysis marked relationships among the major pathways such as negative regulation of wound healing and NABA Extracellular matrix (ECM) (Naba et al., 2012)^14^ regulators associated with renal fibrosis^15^, ionotropic glutamate receptor pathways associated with podocyte cytoskeleton remodeling^16,17^, adaptive immune response pathways, and complement and coagulation cascade. Gene set enrichment analysis (GSEA) revealed significant enrichment of genes associated with adaptive immune response and lymphocyte-mediated immunity in the FSGS group (Fig. 1e, f, Supplementary Table 7c).

### Identification of FSGS-specific protein signature

To further identify specific signature of FSGS, we compared the proteome profiles with those of two disease control groups, MCD and IgAN, using Over representative analysis (ORA) and GSEA (Extended Data Fig. 2a-d, Supplementary Table 8a-9c). Fig. 2a depicts a heatmap of the proteins that varied between the groups. GO pathway analysis showed that complement activation and immune response were enriched in the FSGS group compared with the kidney donor and MCD groups, but not with the IgAN group, in which the complement system actively contributes to the pathogenesis (Fig. 2b).

**Fig. 2.**
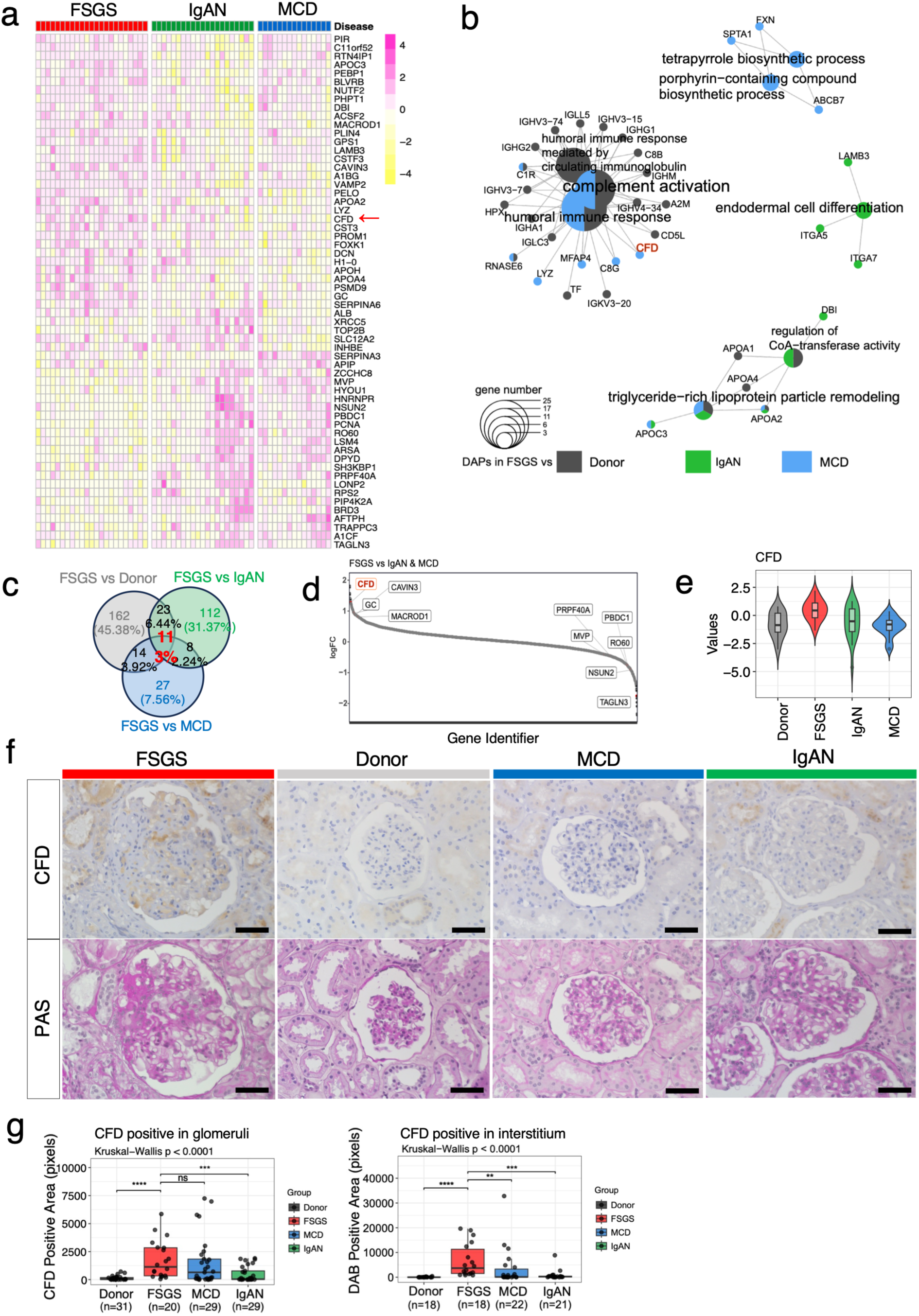
Proteomics analyses comparing the FSGS and disease control groups identified FSGS-specific protein signatures. **a**, Heatmap of top differentially abundant proteins across FSGS, IgAN, and MCD. The top 25 proteins from each comparison (FSGS vs IgAN, FSGS vs MCD, FSGS vs IgAN + MCD), ranked by limma moderated t-statistic, are shown. Color represents row-scaled z-scores of log₂-transformed abundance. **b,** Gene Ontology pathway enrichment analysis of proteins significantly increased in the FSGS group compared with kidney donors, MCD, and IgAN. **c,** A Venn diagram showing the number of significantly increased proteins in the FSGS group compared with kidney donors, MCD and IgAN. Eleven proteins were in common across all the comparisons. The thresholds were |log2 fold change| > 1 for all, and adjusted P < 0.1 for FSGS versus kidney donors and FSGS versus IgAN, and P < 0.1 for FSGS versus MCD. **d,** A rank intensity plot for the panel’s analysis comparing FSGS with two disease control groups. **e,** A box plot displaying protein abundance of CFD in the four groups. **f,** Representative immunohistochemical staining for CFD (top) and PAS staining (bottom) for the four groups. scale bars, 50 µm. **g,** CFD-positive area quantified in glomeruli (left) and tubulointerstitium (right). DAB-positive areas (pixels) per glomerulus or per fixed interstitium area are plotted; n indicates the number of glomeruli or tubulointerstitial regions analyzed for each group. Kruskal–Wallis test with Dunn’s post hoc test (*P < 0.05, **P < 0.01, ***P < 0.001).

There are fewer DAPs between the FSGS and MCD groups (7.56%) when compared to either the kidney donor group (45.38%) or the IgAN group (31.37%) (Fig. 2c); which may reflect substantial overlaps in molecular mechanisms between FSGS and MCD^18^. Venn diagram analysis revealed 11 overlapping proteins that were more than 2-fold enriched in the FSGS group compared to the kidney donor, MCD, and IgAN groups: alpha-1B-glycoprotein, apolipoprotein A-II, apolipoprotein C-III, ADP-ribosylation factor-like protein 6-interacting protein 1, complement C1r subcomponent, complement factor D (CFD), casein kinase II subunit beta, chondroitin sulfate proteoglycan 4, integrin alpha-7, matrilysin, isoform 12 of sorbin and SH3 domain-containing protein 2 (Fig. 2c, Extended Data Fig. 3). Among them, CFD, which activates the alternative complement pathway, exhibited dominant fold changes in the FSGS group in all comparisons against the MCD, IgAN, and combined MCD and IgAN groups (Fig. 2d, 2e, Extended Data Fig. 2e, f). To verify the results of the proteome analysis and determine the sites of CFD accumulation within the kidney tissues, we performed immunohistochemical analysis on the same subjects prepared for LC-MS/MS analysis. In the FSGS specimens, CFD was stained positively in both glomeruli and renal tubules. In contrast, no positive signal was detected in the kidney donor tissues and slight signals were observed in the MCD or IgAN kidney tissues (Fig. 2f, 2g). Collectively, the proteomics and histological analyses demonstrate that immune and complement activation pathways are defining molecular feature of FSGS, with CFD emerging as a potential disease-specific marker.

### Spatially resolved transcriptomics highlights upregulation of complement activating genes in FSGS

Based on these proteomics analyses, we next performed spatial transcriptomics using Xenium in situ on human kidney biopsy specimens from 21 distinct individuals. A custom 100-gene panel was designed to capture molecular events in kidney tissues including complement regulatory genes, reflecting the pathways identified in the proteomics analysis (see Methods and Supplementary Table 10). A total of 477 genes, comprising these 100 genes and 377 genes from the Xenium Human Multi-Tissue and Cancer Panel (1000626; 10x Genomics), were profiled.

Xenium-based cell segmentation that integrated nuclear, cytoplasmic and cell-boundary information enabled identification of glomerular and interstitial structures at single-cell resolution. After quality control and exclusion of cells lacking annotation, 558,976 of the initial 807,440 cells were retained. Dimensionality reduction with Principal Component Analysis (PCA) and Harmony^19^ effectively aligned the obtained datasets of gene expression and minimized the batch effects (Extended Data Fig. 4a). As Robust Cell Type Decomposition (RCTD)-based annotation^20^ mislocalized several glomerular cell types, including parietal epithelial cells (PECs) and interstitial cells. We excluded “PECs” located outside glomeruli. Reclustering “PECs” within glomeruli identified one cluster expressing a mesangial marker, integrin subunit alpha 8 (ITGA8) (Extended Data Fig. 4b, c) and localizing in the mesangial regions (Extended Data Fig. 4d), enabling us to reclassify it as Mesangium 1. The remaining cells expressing a PEC marker, claudin-1 (CLDN1), were annotated as PECs (Extended Data Fig. 4e).

Similarly, intraglomerular cells within the populations initially annotated as “interstitial cells” expressing ITGA8 and localizing in the mesangial regions were classified as Mesangium 2. Both Mesangium 1 and Mesangium 2 were categorized into Mesangium cells for further analyses, consistent with their proximity on the Uniform Manifold Approximation and Projection (UMAP) (Extended Data Fig. 4f). Immune cell annotations were refined through quality control steps to remove cells exhibiting non-immune features or expressing markers indicative of doublets or misannotation. Details on monocyte subpopulation annotation are provided in Extended Data Fig. 4g. Following cell redefinition, 493,066 cells were included in the analysis.

Reclustering integrating spatial coordinates with transcripts refined the characterization of kidney spatial transcriptome architecture (Fig. 3a), with cells appropriately expressing their respective gene markers (Fig. 3b) and localizing inside or outside the glomeruli (Fig. 3c). Cellular composition of the kidney tissues differed between the FSGS and donor groups, with reduced podocytes and proximal tubular cells, increased interstitial cells, as well as immune cells represented by M2 macrophages, and conventional dendric cells (cDC) in the FSGS group (Fig. 3d, e).

**Fig. 3.**
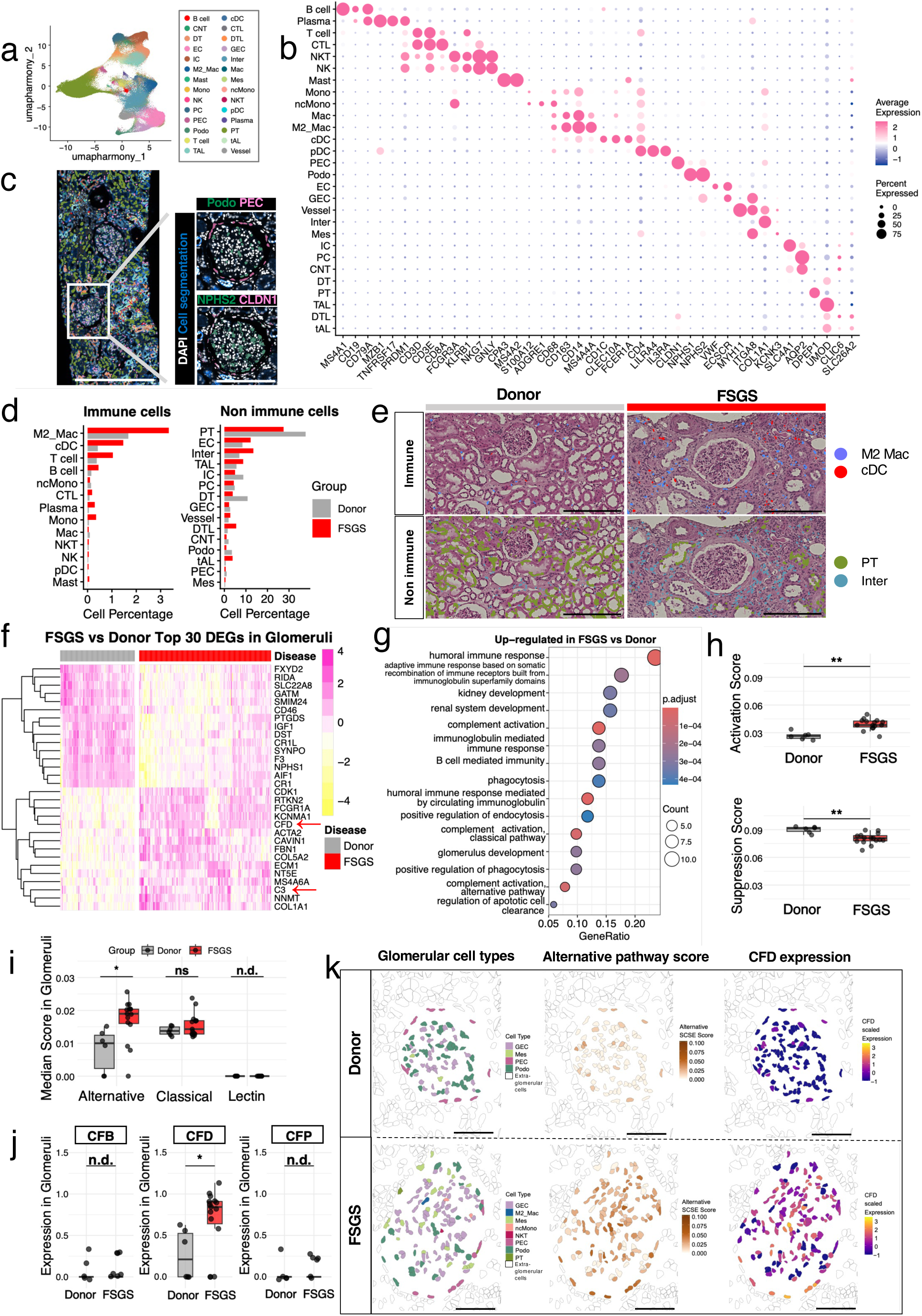
Spatial transcriptomics reveals alternative pathway dominant complement pathway activation in glomeruli of the FSGS group. **a**, UMAP of cell clusters identified by Xenium spatial transcriptomics. Cell types are colour-coded. **b,** A dot plot showing average expression and the percentage of cells expressing cell type specific marker genes across clusters. Colour intensity indicates mean expression, and dot size indicates the percentage of expressing cells. **c,** Representative spatial transcriptomic maps of a kidney tissue section. NPHS2 (podocin) and CLDN1 (claudin 1) mark podocytes and PECs, respectively. Scale bars, left, 500 µm; right, 200 µm. **d,** Percentages of immune (left) and non-immune (right) cell types in the kidney donor and FSGS groups. **e,** Representative images exhibiting the spatial distribution of immune (top) and non-immune (bottom) cells in the kidney donor and FSGS specimens. Scale bar, 250 µm. **f,** A heatmap of DEG in the kidney donor and FSGS specimens. Colour scale indicate normalized expression levels. **g.** GO enrichment analysis of pathways upregulated in glomeruli in the FSGS group. Dot size indicates gene count and colour indicates adjusted p value. **h,i,** Complement scores in the kidney donor and FSGS groups. **h.** Complement activation (top) and suppression (bottom) scores. **i,** SCSE scores for complement initiation pathway of the alternative, classical, and lectin pathways in the glomeruli. Each dot represents the median SCSE score per sample. **j,** Expression of genes associated with alternative pathway activation in the FSGS and kidney donor groups. **k,** Spatial mapping of alternative pathway scores and CFD expression at the single-cell level in the kidney donor and FSGS specimens. Scale bars, 100 µm. *P < 0.05; **P < 0.01; NS, not significant. Cell type abbreviations: DTL, thin descending limb; IC, intercalated cell; PT, proximal tubule; Inter, interstitial cells; CNT, connecting tubule; PC, principal cell; TAL, thick ascending limb; DT, distal tubule; cDC, conventional dendritic cell; tAL, thin ascending limb; PEC, parietal epithelial cell; Podo, podocyte; GEC, glomerular endothelial cell; CTL, cytotoxic T cell; ncMono, non-classical monocyte; M2_Mac, M2 macrophage; EC, extraglomerular endothelial cell; Mes, mesangium; Mono, monocyte; Mac, macrophage; pDC, plasmacytoid dendritic cell

To investigate differences of transcripts in the glomeruli, we defined a glomerulus as one region of interest (ROI) (Extended Data Fig. 4h). Comparative analysis of glomeruli from the FSGS and kidney donor groups revealed increased transcript levels of genes including CFD and C3, alongside reduced podocyte markers including nephrin (NPHS1) and SYNPO (Fig. 3f). Enrichment analysis identified pathways related to humoral immune response, kidney development, and complement activation in the FSGS group (Fig. 3g). Given the data highlighting complement activation as an important feature of FSGS, we performed further analyses focusing on complement-related transcriptional regulation using Single-cell Signature Explorer (SCSE)-based scoring^21^, which quantifies the expression of curated activation– and suppression-related gene sets. The glomerular cells in the FSGS group exhibited significantly higher activation-related and lower inhibitory gene expression scores than those in the kidney donor group (Fig. 3h), which presumably exhibits activated complement-related signals. Scoring of gene sets representing three complement initiation pathways revealed preferential mRNA expression of alternative pathway (Fig. 3i). Among these, CFD was significantly upregulated in the FSGS group (Fig. 3j), consistent with the proteomics analyses. Spatial mapping further demonstrated higher alternative pathway scores and upregulation of CFD mRNA in the FSGS group compared with the donor group (Fig. 3k). These findings collectively highlight activation of the complement alternative pathway at a glomerular-cell level as a unique feature of FSGS that corroborates our proteomics findings.

### Trajectory analyses identify molecular dynamics of podocytes and PECs associated with glomerulosclerosis

Considering the focal and heterogeneous nature of FSGS, we next aimed to examine how these transcriptional alterations at a glomerular level related to disease progression. Setting each glomerulus as a transcriptome unit, we performed trajectory analysis on a diffusion map^22^ originating from the kidney donor dominant cluster using slingshot^23^ (Fig. 4a). To explore glomerular alterations along this trajectory, we divided the pseudotime into quartiles (Extended Data Fig. 5a, b). In the FSGS group, glomeruli were scattered across the quartiles, representing focal progression of FSGS (Fig. 4b) and proportions of podocytes in the glomeruli progressively declined, which coincided with histopathological findings of glomerulosclerosis (Fig. 4c, d), reflecting the course of disease progression.

**Fig. 4.**
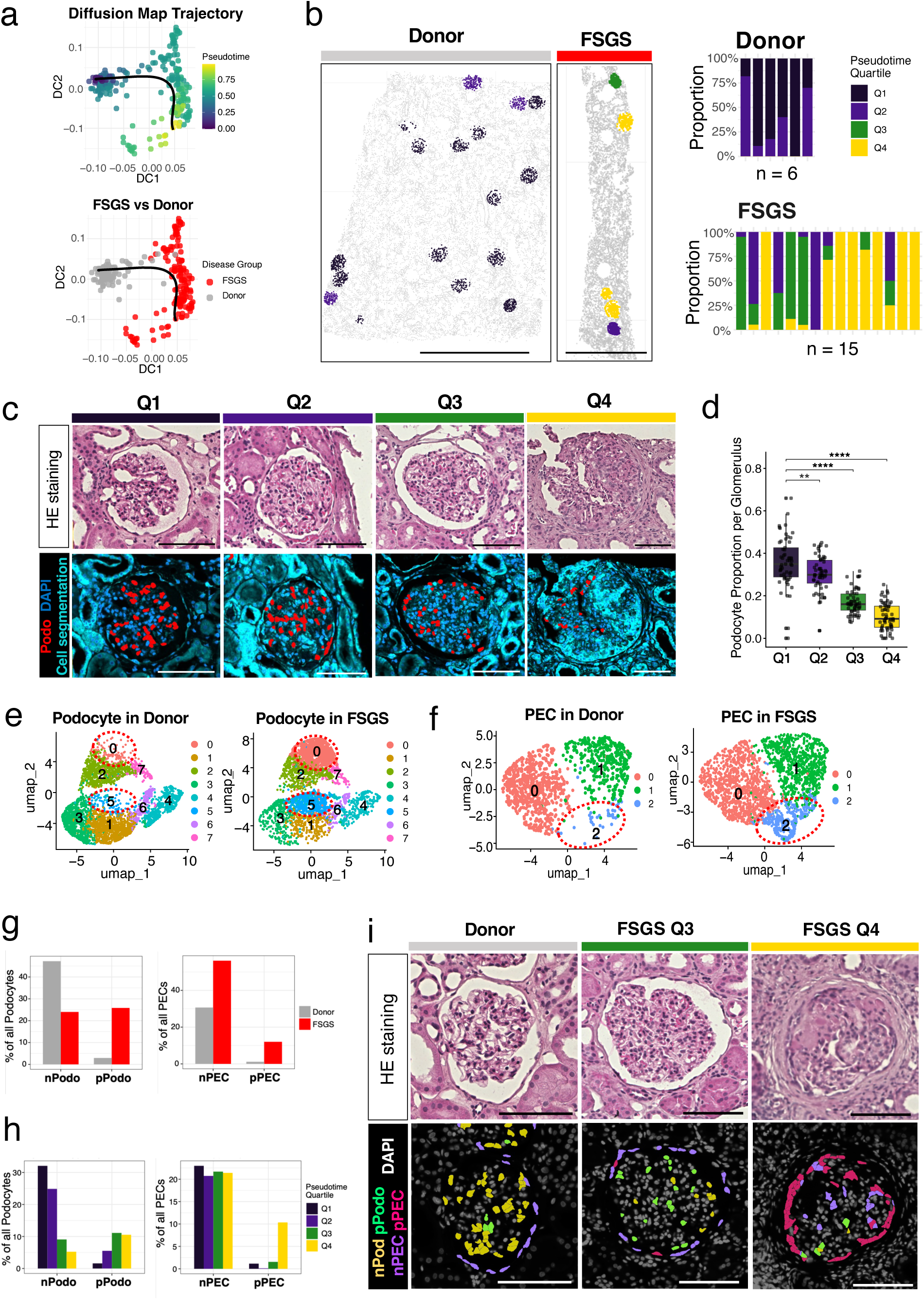
Trajectory analysis reveals progressive podocytes injury and PECs activation in FSGS. **a**, Diffusion map visualization of glomeruli by pseudotime values (top) and disease states (bottom), showing a trajectory along a glomerular injury axis. **b,** Bar plots showing the sample distribution across glomerular quartiles (Q1-Q4, right) and spatial mapping of glomeruli (left) in the kidney donor and FSGS specimens, with individual glomeruli colour-coded by the pseudotime quartile. Scale bars, 1000μm. **c,** Representative HE staining (top row) and spatial distribution of podocytes (bottom row) in the glomeruli belonging to each pseudo-time quartile. Red-coloured podocytes decrease along pseudotime progression. Scale bars, 100μm. **d,** Proportions of podocytes per glomerulus across the pseudotime quartiles. Statistical comparisons were performed using Wilcoxon rank-sum test with Q1 as the reference group. P values were adjusted for multiple comparisons using Bonferroni correction. ****P < 0.0001; ns, not significant. **e, f,** UMAP plots showing clusters of podocytes (e) and PECs (f) in the kidney donor (left) and FSGS (right) specimens, respectively. The red dotted circles (cluster 0 and 5 for podocytes and 2 for PECs) indicate clusters observed predominantly in the FSGS group, which were designated as pathogenic clusters, and the others were defined as normal clusters. **g, h,** Bar plots and spatial mapping showing the distribution of the normal and pathogenic clusters in the kidney donor and FSGS specimens. **i,** Representative HE staining (top row) and spatial distribution of nPodo, pPodo, nPEC, and pPEC (bottom row) in the glomeruli in the kidney donor (left) and FSGS (middle, Q3; right, Q4) specimens. Scale bars, 100μm.

We next examined cellular profiles to identify clusters enriched in the FSGS group. Given the pivotal roles of podocytes and PECs in the pathogenesis of FSGS^24,25,26^, we performed unsupervised clustering followed by UMAP visualization on these two cell types. This analysis identified eight and three clusters in podocytes and PECs, respectively (Fig. 4 e, f). The podocyte clusters, 0 and 5 (Fig. 4e, g), which were predominantly observed in the FSGS group, and were therefore defined as pathogenic podocytes (pPodo). The remaining clusters (1–4 and 6–7) were defined as normal podocytes (nPodo). Similarly, the PEC cluster 2, predominantly observed in the FSGS group (Fig. 4f, g), was defined as pathogenic PEC (pPEC), with the other clusters as normal PEC (nPEC). Along the pseudotime trajectory, proportion of pPodo increased, peaking in Q3, whereas nPodo decreased (Fig. 4h, i). In contrast, nPECs were consistently detected through the pseudotime trajectory, and pPECs were most abundant in Q4 (Fig. 4h), corresponding to glomerular sclerosis observed at an advanced stage (Fig. 4i). These results imply that malfunctions of podocytes and PECs contribute to distinct disease stages during the progression of FSGS.

### Altered intercellular communications centered on podocytes and PECs during FSGS progression

Given the observations that complement signals were activated at a cellular level in the FSGS group, we next assessed complement-related transcriptional changes along the pseudotime. Transcriptional signatures suggestive of complement activation emerged during the glomerular injury, characterized by increasing complement activation scores and declining suppression scores along the pseudotime (Fig. 5a, b). Activation of complement-related pathway appeared to be predominant in podocytes and PECs as both complement activation and suppression scores differed significantly between the FSGS and kidney donor groups, while other glomerular cell types (Extended Data Fig. 6) showed significant changes only in activation scores (Fig. 5c, d).

**Fig. 5.**
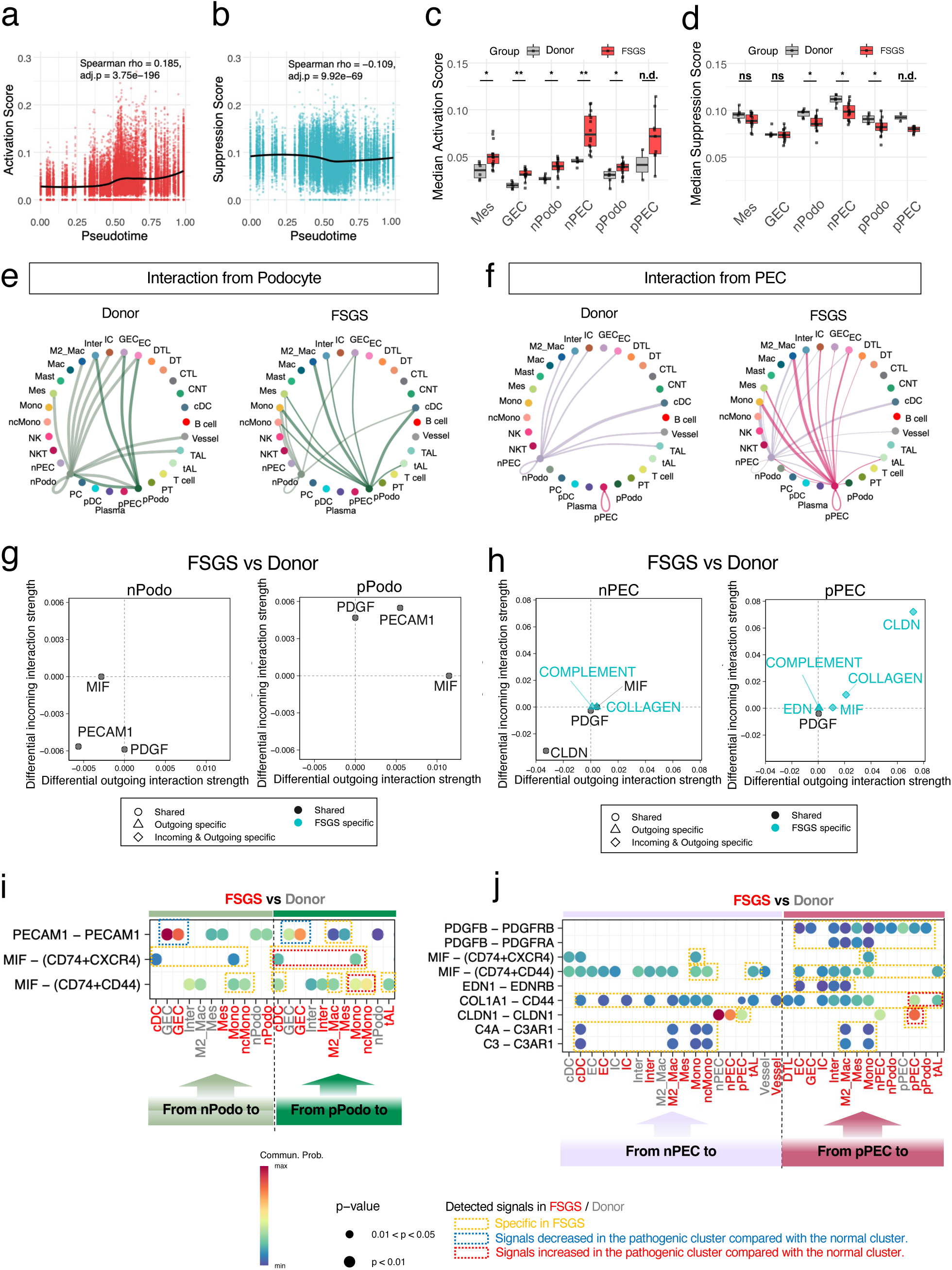
Cell-cell interaction analysis reveals altered intercellular communication in FSGS. **a, b**, Complement activation (a) and suppression (b) scores across the pseudotime quartiles (Q1–Q4). **c, d,** Complement activation (c) and suppression (d) scores across glomerular cell types in the kidney donor (gray) and FSGS (red) specimens. Statistical comparisons were performed using the Mann–Whitney U test with Benjamini–Hochberg correction. *P < 0.05, **P < 0.01; ns, not significant; n.d., not determined (<3 samples per group). **e, f,** Network visualization of podocyte– (e) and PEC-originating (f) cellular interactions in the kidney donor (left) and FSGS (right) specimens. Lines are coloured as follows: light yellow-green, nPodo; dark green, pPodo; magenta, pPEC; light purple, nPEC. Line thickness indicates interaction strength. **g, h,** Differential interaction strength analyses for normal/pathogenic podocytes (g) and PECs (h) comparing the FSGS with kidney donor groups. Incoming (y axis) and outgoing (x axis) interactions are shown with symbol types indicating interaction categories. Gray indicates interactions observed in both donor and FSGS groups, while light blue indicates FSGS-specific interactions. Shapes represent interaction directionality: triangles, outgoing-specific; diamonds, incoming-specific; circles, observed in both directions. **i, j.** Bubble plots of ligand–receptor interactions in podocytes (i) and PECs (j) comparing the FSGS and kidney donor groups. Dot size indicates P values; dot colour gradient indicates communication probability. Dashed squares in yellow, blue, and red indicate signals specifically detected in FSGS, signals decreased in FSGS, and signals increased in FSGS, respectively. Abbreviations: MIF, macrophage migration inhibitory factor; PECAM1, platelet endothelial cell adhesion molecule 1; PDGF, platelet-derived growth factor; CLDN, claudin; EDN, endothelin.

To investigate molecular events in podocytes and PECs including the clusters (pPod and pPEC) increased over the pseudotime, we analyzed spatially constrained intercellular communications of these cells using CellChat^27^. In the analysis, both podocytes and PECs, particularly pPodo and pPEC, exhibited increased intercellular interactions in the FSGS group compared with the kidney donor group (Fig. 5e, f). Podocytes exhibited reinforced connections with immune cells, including monocytes, cDC, and M2 macrophages, while PECs showed interactions shifted from homotypic to heterotypic with cells involving immune cells such as monocytes and M2 macrophages, podocytes and tubular cells (Fig. 5f).

To further characterize these altered signaling patterns, we performed ligand– receptor analysis. pPodo exhibited increased platelet-derived growth factor (PDGF), platelet endothelial cell adhesion molecule 1 (PECAM1), and macrophage migration inhibitory factor (MIF) signaling (Fig. 5g). By contrast, PECs displayed several pathways such as collagen and complement, predominant in the FSGS group at the early stage, and claudin (CLDN), endothelin (EDN) and MIF signaling emerging as injury progressed (Fig. 5h). In the FSGS group, PECAM1–PECAM1 signaling from podocytes to glomerular endothelial cells (GECs) was markedly reduced in pPodo compared with nPodo at the late stage (Fig. 5i), suggesting impaired podocyte–GEC interactions that are critical for the maintenance of podocyte function^28^. Podocytes also demonstrated enhanced MIF signaling via CD74–CXCR4/CD44 interactions with immune cells such as monocytes, dendritic cells, M2 macrophages and tubular cells in the FSGS group (Fig. 5i), consistent with previous reports that MIF signaling contributes to podocyte injury and inflammatory activation in glomerular diseases^29,30^.

PECs in the FSGS group exhibited abundant PDGF signaling, involving both interactions homotypic and heterotypic to cells inside and/or outside the glomeruli (Fig. 5j), suggesting that this enhanced signaling may lead to PEC activation and increased extracellular matrix, as reported in previous studies^31,32,33^. Moreover, PECs showed several signaling pathways augmented in the FSGS group; CD74-CXCR4/CD44 mediated MIF signaling with endothelial, mesangial cells, immune cells and tubular cells, EDN1–EDNRB signaling, COL1A1–CD44 interactions with thin ascending limb cells, CLDN1–CLDN1 signaling among pPEC cells and complement ligand–receptor (C3/C4A–C3aR1) interactions with M2 macrophages and monocytes (Fig. 5j), suggesting that pPECs acquire adhesive and proinflammatory phenotypes, which contribute to progression of glomerular inflammation and fibrosis.

Collectively, spatial analysis revealed coordinated remodeling of cellular communication networks in glomeruli of FSGS, in parallel with complement-related signal activation.

### Spatial propagation of complement-related signals emanates from injured glomeruli

Complement activation scores were also elevated in interstitial regions in the FSGS group (Fig. 6a), which prompted us to investigate spatial and functional relationships between intra– and extra-glomerular molecular events. To address this, we classified interstitial regions into distance-based quartile categories from glomeruli: Very Close (0-25th percentile of the distance from glomeruli), Close (25-50th percentile), (50-75th percentile), and Far (75-100th percentile) (Fig. 6b, c, Extended Data Fig. 7). Cells in the Very Close regions exhibited complement activation scores increased depending on the progression of glomerular injury, whereas no significant difference was observed in those in the Far regions (Fig. 6d). Based on these results, we further focused our analysis on the Very Close regions and observed higher complement activation scores at advanced (Q4) than early (Q1) stages of FSGS around glomeruli (Fig. 6e-g), suggesting a spatial propagation of complement activity from injured glomerular cells to those in the neighboring interstitial regions.

**Fig. 6.**
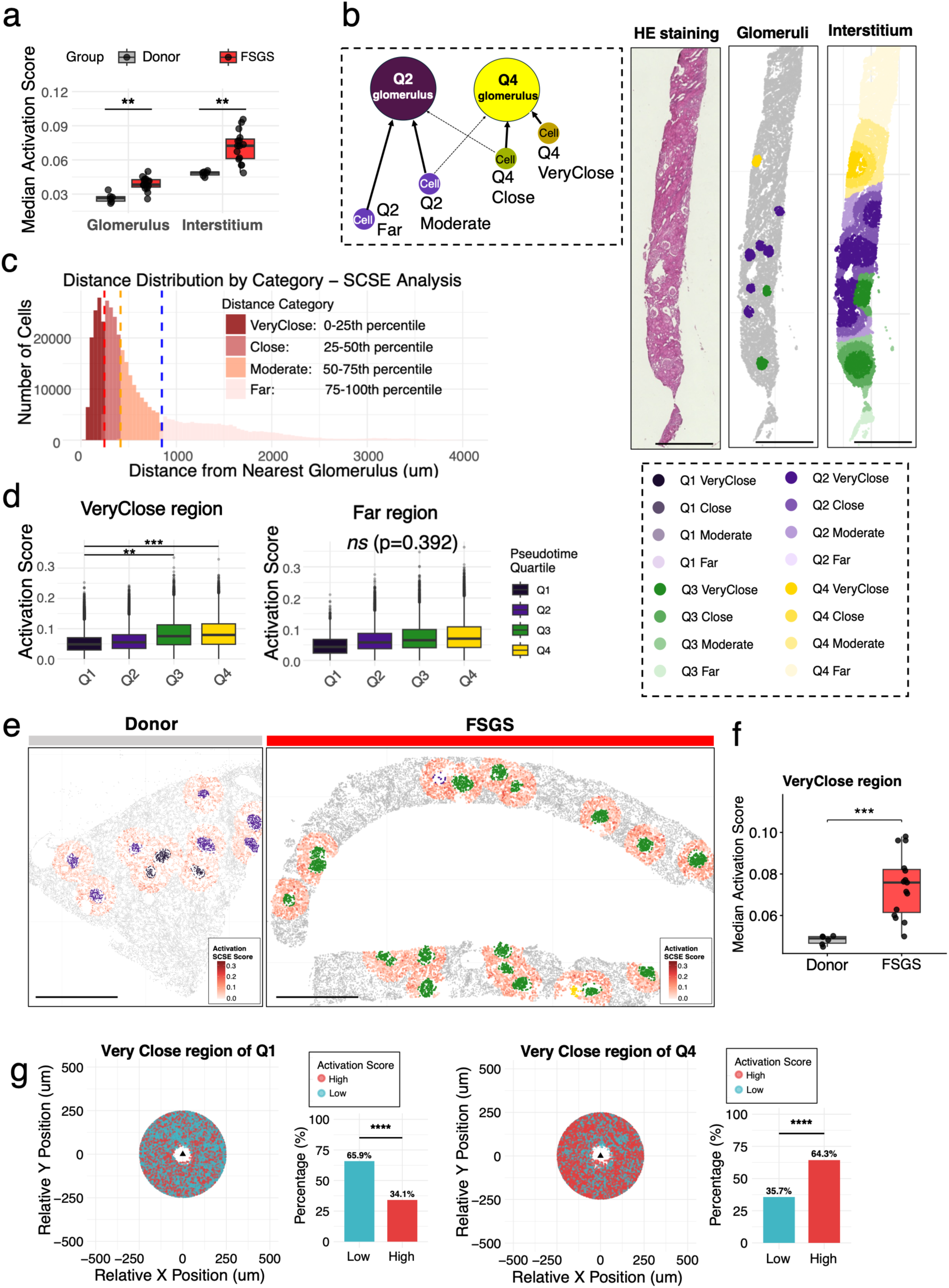
Spatial gradient of complement activation from injured glomeruli. **a**, Median complement activation SCSE scores for glomeruli and interstitium in the kidney donor (gray) and FSGS (red) groups. *P < 0.05, **P < 0.01, ***P < 0.001 (Mann-Whitney U test). **b,c,** Schematic figure (left) and representative spatial mapping (right) of the kidney tissue sections classified by the pseudotime quartiles (Q1-Q4) and the distance quartiles from glomeruli (c, Very Close, 0-25th percentile of the distance from glomeruli, 0-246.26μm; Close (25-50th percentile, 246.26-415.31μm); Moderate (50-75th percentile, 415.31-847.63μm); Far (75-100th percentile, 847.63-4043.31μm). Scale bar, 1000 μm. **d,** Complement activation SCSE scores of the interstitial cells in the Very Close region (left) and the Far region (right) across the pseudotime quartiles. **e, f,** Spatial distribution (left) and quantification (right) of complement activation SCSE scores in the Very Close region for the kidney donor and FSGS groups. **g,** Spatial distribution of complement activation SCSE scores in the Very Close region surrounding the glomerulus for Q1 (left) and Q4 (right), displayed in relative coordinate space centered on the glomerular centroid. Cells with high (red) or low (gray) activation, defined by median-based binarization within each distance category, are shown. The triangular marker indicates the glomerular center and axes represent relative positions from the glomerulus. ****P < 0.0001 (Chi-square test).

### Remodeled intercellular communication networks mediate spatial disease propagation

To explore molecular mechanisms underlying the spatial relationship between glomerular injury and molecular events occurred in adjacent interstitial regions, we examined dynamics of intercellular molecular signals in the glomeruli and the Very Close regions throughout the pseudotime. Analysis of intercellular communication revealed that the overall strength of intercellular interactions was highest in Q3 and decreased in Q4, the most advanced stage of glomerulosclerosis (Extended Data Fig. 8a). Infiltrating immune cells increased at Q3 (Fig. 7a), predominantly M2 macrophages and cDCs, accompanied by a reduction in tubular epithelial cells and an expansion of interstitial cells (Fig. 7b), which associates with fibrosis in injured glomeruli at an advanced stage. Signals involving CD45, complement, and C–C motif chemokine ligand (CCL) pathways, which associate with immune cell recruitment and lymphocyte homing^34,35^, as well as collagen pathway, differed significantly between the FSGS and donor groups (Fig. 7c). Among these pathways, only collagen signaling showed a continuous increase up to Q4 with glomerulosclerosis, whereas the other pathways did not show this trend (Fig. 7d).

**Fig. 7.**
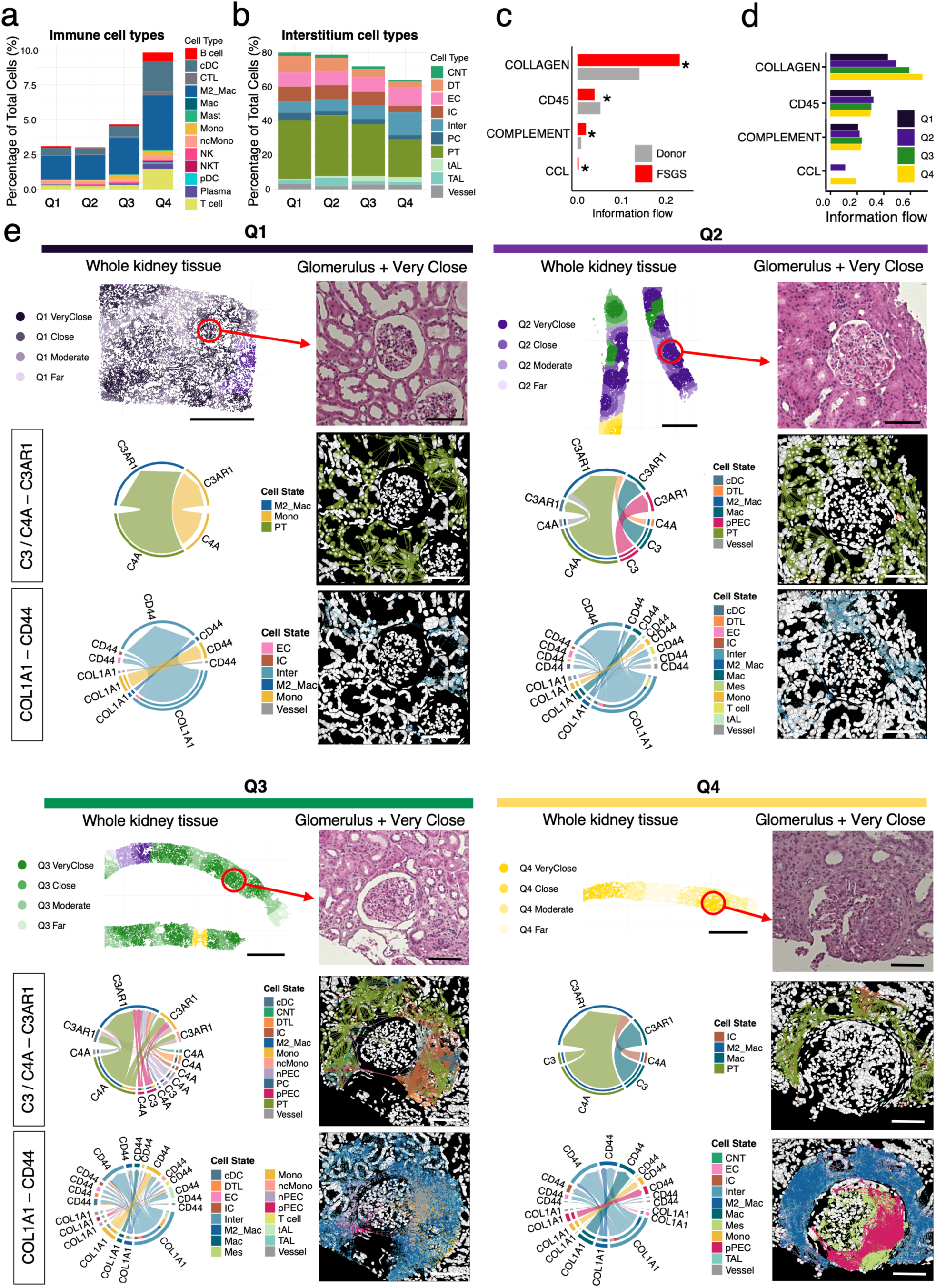
Remodeled intercellular communications driving spatial disease propagation. **a, b**, Composition of immune (a) and interstitial cell types (b) across the pseudotime quartiles in the glomeruli and Very Close region. Stacked bar charts indicate the percentage of subpopulations in total cells. **c,** Signaling pathways significantly altered in the FSGS group (red) compared with kidney donors (gray). **d,** Information flow analysis of the specific signals in the FSGS group shown in panel (c), stratified by glomerular pseudotime quartiles (Q1–Q4). **e,** Visualization of ligand-receptor interactions in the glomeruli and Very Close regions across all glomerular quartiles. Upper panels are images of whole kidney tissues belonging to the pseudotime quartiles. The glomeruli and Very Close regions are magnified and shown as HE stainings. Middle and lower panels show chord plots depicting cell-cell communication networks via C3/C4A-C3AR1 and COL1A1-CD44 axes and their spatial visualization mapped onto representative tissue sections. Scale bar, 1000 μm (top); 100 μm (middle, bottom). Cell type abbreviations: DTL, thin descending limb; IC, intercalated cell; PT, proximal tubule; Inter, interstitial cells; CNT, connecting tubule; PC, principal cell; TAL, thick ascending limb; DT, distal tubule; cDC, conventional dendritic cell; tAL, thin ascending limb; nPEC, normal parietal epithelial cell; pPEC, pathogenic parietal epithelial cell; nPodo, normal podocyte; pPodo, pathogenic podocyte; GEC, glomerular endothelial cell; CTL, cytotoxic T cell; ncMono, non-classical monocyte; M2_Mac, M2 macrophage; EC, extraglomerular endothelial cell; Mes, mesangium; Mono, monocyte; Mac, macrophage; pDC, plasmacytoid dendritic cell

We then sought to identify molecular pathways that capture these stage-dependent changes, focusing on the complement pathway, which was prominent in both our transcriptomics and proteomics analyses, and the collagen pathway, which plays important roles in renal fibrosis^36^. In the FSGS group compared to the kidney donor group, complement signaling closely associated with interactions between PECs and M2 macrophages whereas enhanced collagen signaling involved complex intercellular networks among various cell types including PECs, interstitial cells, macrophages, tubular cells and endothelial cells (Extended Data Fig. 8b). We next analyzed cellular interactions across Q1 to Q4, allowing us to map the dynamics of immune and fibrotic processes along the pseudotime. This analysis captured C3/C4A–C3AR1 and COL1A1– CD44 axes represented of the complement-related and collagen-related interactions, respectively (Fig. 7e). Intracellular complement activation signal initially arose from proximal tubular epithelial cells and monocytes toward M2 macrophages at Q1, expanded at Q2 to include pPECs, cDCs, vascular cells, and distal tubule cells, and involved the broadest range of interacting cell types at Q3 with additional participation of nPECs and interstitial cells. Complement interactions were largely reduced at Q4 and confined mainly to tubular and macrophage populations. In contrast, collagen signaling, driven primarily by interstitial cells at Q1 with involvement of monocytes, progressively incorporated cDCs, mesangial cells, distal tubule cells at Q2, and nPECs and pPECs at Q3, and persisted up to Q4 via interstitial cells, pPECs, and mesangial cells, reflecting sustained fibrosis and ongoing tissue remodeling.

These results, collectively, reveal the dynamic of intercellular communications among glomerular, interstitial and infiltrating immune cells such as macrophages and cDCs, which drives pathogenic molecular signals originated from glomeruli to whole kidney tissues during disease progression.

## Discussion

In this study, we applied comprehensive proteomics and spatially resolved transcriptomics to delineate the molecular and cellular architecture of kidney tissues of FSGS. We provide a high-depth, unbiased proteomics dataset of human kidney biopsy tissues, alongside spatial transcriptomics data generated using the Xenium in situ platform, which achieves single-cell resolution. By examining multi-layered omics data at protein and mRNA levels, we captured a comprehensive view of molecular events underlying the pathogenesis of FSGS. Glomerulus-oriented trajectory analysis enabled us to identify dynamics of cell type–specific signaling during disease progression. Collectively, our multi-layered omics data provide a valuable resource and lay the groundwork for future glomerular studies.

Building on our proteomics results that complement activation is prominently enriched in the FSGS group, we examined spatial influences of this dysregulated complement activity at single-cell resolution on disease progression. Unsupervised clustering followed by UMAP visualization identified FSGS-enriched clusters predominantly observed within podocytes and PECs populations (Fig. 4e, f). Intercellular interactions originated from podocytes and PECs altered as glomerular injury progressed. Damaged podocytes showed reduced PECAM1 signaling and increased MIF signaling (Fig. 5g, i), consistent with prior reports that loss of podocyte– endothelial interactions and elevated MIF contribute to glomerular injury and macrophage/dendritic cell activation^37, 38,39, 40,41^. These observations are consistent with our interpretation that dysregulated podocyte-mediated crosstalk may act as an early driver of inflammatory amplification in FSGS. We further identified PDGF, CLDN1, EDN, MIF, collagen, and complement signaling derived from PECs, contributing to the pathogenesis of FSGS (Fig. 5h, j). These signatures represent activated PECs, which is implicated in promoting collagen deposition and glomerulosclerosis in FSGS and other glomerulonephritides^24,25,36,42,43,44,45,46^, confirming that injured podocytes and PECs play central roles in inflammatory aspects of FSGS pathology.

Our proteomics profiling revealed immune and complement pathways as central components of molecular landscapes in FSGS (Fig. 2b–d). These findings align with accumulating evidence implicating immune dysregulation^8,47,48,49,50^ and complement abnormalities via both the classical and alternative pathways^49,51,52,53,54,55,56,57,58,59^ in the pathogenesis of FSGS. Importantly, we identified CFD as a specific marker for FSGS. Spatial transcriptomics further validated increased CFD expression at the mRNA level and demonstrated preferential activation of the alternative complement pathway within glomeruli, suggesting that CFD may also serve as a driver of complement-mediated injury in FSGS. This notion is corroborated by previous experimental findings in the adriamycin induced FSGS mouse model, where CFD deficiency reduced proteinuria and attenuated structural damage^54^. These findings together support the concept that targeting CFD may offer a potential therapeutic strategy for human FSGS.

Emerging data highlight the importance of intracellular complement signaling, known as complosome, which regulates cellular metabolism, mitochondrial activity, survival, and gene regulation in both immune and non-immune cells^60,61^. Intracellular complement activity has been observed in several kidney cell types and is now recognized as a part of intrinsic responses against kidney injury. Previous study showed contribution of complosome to the pathogenesis of FSGS, by revealing increased C5aR1 expression in kidney tissues of primary FSGS patients and beneficial effects of its blockade in animal FSGS models^62^. In our study, podocytes and PECs in the FSGS group exhibited elevated complement gene signatures. Previous reports show that podocytes, which express complement components^63,64,65^, contribute to local glomerular complement activation and regulation^64,66^. Human PECs constitutively express C3aR, and exposure to C3a in vitro enhances PEC proliferation and CXCR4-associated migration, supporting a role for C3/C3a in PEC activation in proteinuric nephropathy^67^. Furthermore, in kidney tissues from patients with glomerulopathies, glomerular deposition of C3/C3a parallels PEC activation and upregulation of CXCR4 and related markers^67^. Our data in this study show that PECs participate in delivering complement related signals to monocytes, macrophages, and cDC (Fig. 5j). Macrophages play roles in complement-mediated responses^68^, as reported that increased local C3, which is produced mainly from monocytes/macrophages, drives renal fibrosis^69^. In addition, anaphylatoxins C3a and C5a have been reported to modulate the function of human monocyte-derived DCs^70^, suggesting that complement signals delivered by PECs may also influence cDC immunoregulatory functions.

In addition to complement-mediated interactions, our spatial ligand–receptor analyses revealed that pathogenic podocytes and PECs engage distinct signaling programs that may further amplify inflammatory and fibrotic responses. pPodo exhibited enhanced PDGF, PECAM1, and MIF signaling, together with MIF signaling via CD74–CXCR4/CD44 interactions, which have been implicated in macrophage recruitment, immune cell activation, and podocyte injury in glomerular diseases^29,30,31,32,33,37, 38,39, 40,41^. In parallel, pPECs acquired adhesive and proinflammatory phenotypes characterized by augmented PDGF, MIF, collagen, and cell–cell adhesion signaling, consistent with previous reports linking activated PECs to extracellular matrix deposition, immune cell interactions, and progression of glomerulosclerosis^31,32,33,42,43,44,36,45,46^. Together, these findings suggest that altered podocyte and PEC signaling programs may cooperate to sustain inflammatory amplification and fibrotic remodeling in FSGS. Collectively, PEC-derived complement signals may act to prime macrophages and other immune cells, which may contribute to initiation and progression of glomerulosclerosis in FSGS.

Certain aspects of our study should be taken into accounts. The analyses were conducted on a limited set of human kidney biopsy samples, which provide a representative snapshot of FSGS but may not capture the full spectrum of disease heterogeneity. In addition, the targeted gene panel used for spatial transcriptomics covers only a subset of relevant genes, so some additional cell type–specific interactions might emerge in future studies.

Importantly, these multi-layered omics datasets offer high-resolution resources for understanding pathogenesis of FSGS and facilitating further mechanistic investigations. Our findings altogether delineate distinct molecular stages in FSGS as intracellular complement signals, primarily centered on podocytes and PECs in glomeruli, result in recruitments of immune cells and subsequent fibrosis, propagating into whole kidney tissues, which may benefit for developing stage-specific therapeutic approaches.

## Data Availability Statements

The mass spectrometry and resulting search data have been deposited to the ProteomeXchange Consortium via the PRIDE^71^ partner repository with the dataset identifier PXD052086 (ref: 10.6019/PXD052086). The analysis code and associated data for the proteomics and Xenium spatial transcriptomics analyses have been deposited in Figshare (https://doi.org/10.6084/m9.figshare. [10.6084/m9.figshare.31316515]) and will be made publicly available upon publication.

## Acknowledgements

This study was in part supported by the research grants from the American Heart Association (23CDA1052394) to ARS, Kowa Company, Ltd, Nagoya, Japan to MA and Grants-in-Aid for Scientific Research from the Japan Society for the Promotion of Science (21H02950) to YK. We would like to acknowledge the institutions that cooperated in obtaining consent. We also thank the Research Support Center and Kyushu University Graduate School of Medical Sciences for technical support, M. Munakata and M. Tanaka at the Department of Medicine and Clinical Science for histology assistance, and BioRender for graphical abstract creation. We thank Gabrielle White Wolf, PhD, from Edanz (https://jp.edanz.com/ac) for editing a draft of this manuscript.

## Authors’ contributions

S.H, S.Y., T.N. and Y.K. conceptualized the study. S.H. performed LC-MS/MS measurements, proteomics data acquisition, Xenium experiments and spatial transcriptomics data analysis, and immunostaining, and writing the manuscript. M.T. performed Xenium experiments and analysis. D.S. supervised LC-MS/MS measurements and data acquisition. A.R.S. and S.A.S. performed the proteomics data analysis and interpreted data. S.A.S. contributed to data acquisition in Proteome Discoverer. T.I., H.K. and A.T. collected the samples and helped the experiments. T.N., D.S. and Y.K. supervised the whole experiments and edited the manuscript. M.A., D.K., S.Y., T.A. and T.K. reviewed and edited the manuscript.

## SUPPLEMENTARY MATERIAL

### Extended Data Figures

Extended Data Fig. 1. Quality assessment of kidney tissue composition and proteomics data.

Extended Data Fig. 2. Differential protein abundance in FSGS and disease controls.

Extended Data Fig. 3. Eleven proteins elevated in FSGS relative to healthy and disease controls.

Extended Data Fig. 4. Integration, annotation, and refinement of Xenium spatial cell states.

Extended Data Fig. 5. Pseudotime analysis of glomerular cells reveals disease-associated transcriptional trajectories.

Extended Data Fig. 6. Glomerular cell type distribution in FSGS and donors.

Extended Data Fig. 7. Classification of interstitial regions according to glomerular pseudotime and distance.

Extended Data Fig. 8. Cell–cell interaction analyses across disease states and glomerular pseudotime in glomerular and periglomerular regions.

### Supplementary information

#### Method

Supplementary TableS1. The patients background at kidney biopsy in proteomics cohort.

Supplementary Table S2. The patients background at kidney biopsy in spatial transcriptomics cohort.

Supplementary Table S3. Classification, treatment, and electron microscopic findings of FSGS groups used in proteomics and spatial transcriptomics cohorts.

Supplementary Table S4. Evaluation of pathological differences in the kidney biopsy samples among the four groups in proteomics cohort.

Supplementary TableS5. Evaluation of pathological differences in the kidney biopsy samples among the two groups in spatial transcriptomics cohort.

Supplementary TableS6. All groups consensus proteome with quantification (exported from Proteome Discoverer 2.5)

Supplementary TableS7a. FSGS vs Kidney Donor_DAP results

Supplementary TableS7b. FSGS vs Kidney Donor_GO ORA_enriched in FSGS

Supplementary TableS7c. FSGS vs Kidney Donor_GSEA

Supplementary TableS8a. FSGS vs MCD_DAP results

Supplementary TableS8b. FSGS vs MCD _GO ORA_enriched in FSGS

Supplementary TableS8c. FSGS vs MCD _GSEA

Supplementary TableS9a. FSGS vs IgAN_DAP results

Supplementary TableS9b. FSGS vs IgAN _GO ORA_enriched in FSGS

Supplementary TableS9c. FSGS vs IgAN _GSEA

Supplementary TableS10. The list of target and base panel genes

Supplementary TableS11. The list of genes used for SCSE analyses

## METHODS

### Patients, ethical approval, and tissue collection

This study included adult patients who underwent kidney biopsy at Kyushu University Hospital and eight cooperating hospitals between 2011 and 2024. Each participating hospital obtained informed consent using the opt-out method, under the approval of the Ethics Committee of Kyushu University (approval number 22295-01). All kidney biopsy specimens were evaluated according to established criteria for FSGS based on histopathological findings^72^. Following routine histopathological diagnosis, cases with sufficient residual tissue were enrolled. Kidney biopsy specimens were obtained as part of routine diagnostic procedures and were preserved until subsequent experimental use. This study was conducted following the Declaration of Helsinki and Declaration of Istanbul.

### Sample preparation for LC-MS/MS

Six 15-µm-thick sections were prepared from frozen kidney blocks at −20 °C. The presence of glomeruli was confirmed by light microscopy, and three sections were subsequently placed into a 1.5-mL tube. The serial sections cut for proteome analysis were sliced 4 µm thick and stained with hematoxylin-eosin using a BZ-X800 analyzer (KEYENCE, Osaka, Japan), to estimate the percentage of input glomeruli for LC-MS/MS. We extracted proteins from the tissue using the Qproteome® FFPE Tissue Kit (37623; Qiagen, Venlo, Netherlands), without deparaffinization according to the manual. The purified protein precipitate was dissolved in 50 µl of 8 M urea in 500 mM Tris HCl (pH 8.0). The protein concentration was measured with the BCA Protein Assay (23228; Thermo Fisher Scientific, Waltham, MA, USA). After equalizing the protein amount, Trypsin/Lys-C Mix (VA1061; Promega, Madison, WI, USA) was added at 1:10 and the sample was incubated at 70°C for 1 hr for protein digestion following the manufacturer’s instructions. The peptide solution was then desalted and fractionated using GL-Tip^TM^ SDB-SCX (7510-11202; GL Sciences, Tokyo, Japan), and eight fractions were made sequentially following the manual. The first fraction did not give a good chromatogram when applied to LC-MS/MS (data not shown), thus we use the second through the eighth fractions for the measurement. The flow-through of the last fraction was further absorbed using GL-Tip^TM^ GC (7820-11201; GL Sciences). Two adjacent fractions were combined; finally, four fractions per subject were prepared and each fraction was measured by LC-MS/MS.

### LC-MS/MS measurement and data acquisition

We applied each fraction to an Evotip (EV2001; AMR, Inc. Tokyo, Japan) and separated with HPLC (EVOSEP ONE; AMR, Inc.) using a 15 cm × 150 µm of 1.5 µm column (EV1137; AMR, Inc.) heated to 40°C. The column was connected to the Q-Exactive Orbitrap mass analyzer (Thermo Fisher Scientific) through the emitter (EV1111; AMR, Inc.), and each sample was separated for 88 min gradient time at a rate of 0.22 µL/min with manufacturer’s preset gradient conditions. Detailed data acquisition methods are described in Supplementary information. The peptide false discovery rate (FDR) was calculated using Percolator (target/decoy method) and FDR was set to 0.01 for high-confidence peptides. Protein abundance was estimated using the Proteome Discoverer’s non-label quantification algorithm. The data were normalized using the ‘total peptide normalization’, the default/recommended parameter within PD 2.5.

### Normalization and statistical analysis of proteomics data

We further normalized the exported proteome with equal median (EM) normalization using R version 4.2.2. Missing values were replaced with 0.01 and a base-2 logarithmic transformation was performed. Proteins with missing values in more than 89% of samples (>65 of 73) were excluded from downstream analysis. Differential abundance analysis was performed using a linear model with group as a fixed effect (design formula: ∼0 + Group) fitted with the limma package (3.54.2)^73^, which applies moderated t-statistics. A protein was considered differentially abundant if it had |log_2_FC| > 1 and an adjusted *P* values < 0.05 for comparison with the kidney donor group, and nominal *P* value < 0.05 for disease comparisons. We performed over-representational analysis (ORA) for proteins with |log_2_FC| >1 and GSEA using ranked log_2_FC with GO and KEGG database using R (version 4.2.2) package clusterProfiler (version 4.6.2)^74^, and AnnotationDbi R package “org.Hs.eg.db” (version 3.16.0) was used to map gene identifiers. In the GSEA network visualization, similar pathways were clustered using the aPEAR package in R^75^, with circle size representing the number of related pathways. Pathways with FDR < 0.1 were considered significant. For FSGS versus kidney donor, we used multiple pathway databases such as canonical pathways, GO, KEGG, Panther, Reactome, and WikiPathways using Metascape^76^.

### Immunohistochemical staining

We sliced formalin-fixed, paraffin-embedded human kidney tissue samples into 3 µm sections. The samples were deparaffinized, rehydrated, and washed, followed by antigen retrieval (424142; Nichirei Biosciences, Tokyo, Japan). After incubation in 30% H_2_O_2_ in methanol and then Blocking One (03953-95; Nacalai Tesque, Kyoto, Japan), the sample was incubated with primary antibody against complement factor D (MA5-43873; Invitrogen, Thermo Fisher Scientific Inc.) at 4°C overnight. The sample was then incubated with secondary antibody (424142, Nichirei Biosciences) for 30 min at room temperature and then developed with DAB (415171; Nichirei Biosciences).

Glomeruli and predefined tubulointerstitial regions were extracted as individual image files using a BZ-X800 analyzer (KEYENCE, Osaka, Japan). The extracted TIFF images were then analyzed in R using the EBImage^77^. DAB-positive pixels were automatically detected by applying RGB thresholds (red > 0.3, green > 0.3, blue < 0.6), followed by quantification of the DAB-positive area.

### The workflow for Xenium analysis

FFPE tissue was thinned into 5 µm according to the manual (Demonstrated Protocol CG000578; 10x Genomics, Pleasanton, CA, USA). Tissue slides were analyzed on a Xenium analyzer (Analysis version 2.0.0.10 and software version 2.0.1.0; 10x Genomics, Pleasanton, CA, USA) after deparaffinization, decrosslinking, probe hybridization and wash, ligation, amplification, cell segmentation and blocking, stain enhancement, and autofluorescence quenching in accordance with the manuals (Demonstrated Protocol CG000580, CG000749, CG000582, CG000584; 10x Genomics, Pleasanton, CA, USA). After the run, the tissue slides used for Xenium analysis were HE-stained according to protocol (Demonstrated Protocol CG000613; 10x Genomics, Pleasanton, CA, USA), and the HE stained-images and Xenium transcript data were superimposed using Xenium Explorer version 3.2.0.

### Quality control, normalization, and cell annotation of spatial transcriptomics data

Xenium spatial transcriptomics data were analyzed in R (v4.4.2) using Seurat (v5.3.0). Large language models were used in a limited capacity to assist with code debugging and optimization during Xenium spatial transcriptomics data analysis, and all analytical decisions and interpretations were made by the authors. Cell segmentation was performed using the Xenium Cell Segmentation Kit with the default algorithm provided by 10x Genomics based on DAPI nuclear staining, cytoplasmic and cell boundary staining. For quality control, cells with zero transcript counts were excluded, and cells with transcript counts outside the 5th–95th percentile range were removed from each sample to eliminate low-quality cells and potential segmentation artifacts. Duplicate cell IDs identified across datasets were removed prior to merging. Gene expression matrices were normalized using the NormalizeData function with the median transcript count as the scale factor. To account for batch effects across samples, dataset integration was performed using Harmony^19^. PCA was applied for dimensionality reduction (dims = 20). Cell type annotation was performed using RCTD^20^ with doublet mode, excluding cells with fewer than 50 total Unique Molecular Identifier (UMI) counts, and the cell type with the highest deconvolution weight was assigned to each cell. A reference dataset was obtained from single-cell RNA sequencing data of human kidney tissue [https://datasets.cellxgene.cziscience.com/1c360b0b-eb2f-45a3-aba9-056026b39fa5.rds, Downloaded on July 31, 2024], retaining only cell types with at least 25 cells. Cell type annotations were subsequently refined using spatial context and marker gene expression. Cells initially annotated as podocytes, PECs, endothelial cells, and interstitial cells were classified as intraglomerular or extraglomerular based on their localization within or outside segmented glomeruli. Monocytes were reclassified into classical and non-classical subtypes based on established subtype-specific marker expression. Immune cells were further filtered using negative marker-based quality control, removing cells expressing stromal (COL1A1), endothelial (VWF, ECSCR), or lineage-inappropriate markers.

### Expression-based scoring of complement pathways

To evaluate the regulation of the complement system, we manually curated complement-related gene sets from genes retrieved by searching the National Center for Biotechnology Information (NCBI) database with the term “complement”, and calculated enrichment scores using the SCSE method^21^. The SCSE score for each cell was computed as the sum of normalized expression values of genes in the gene set divided by the total expression of all genes in that cell. Gene sets were defined for complement activation (C1QA, C1QB, C1QBP, C1QC, C1R, C1S, C2, C3, C4A, C4B, C5, C6, C7, C8A, C9, CFB, CFD, CFP, FCN2, MASP1, MASP2, and MBL2), complement suppression (C4BPA, C4BPB, CD46, CD55, CD59, CFH, CFHR1–5, CFI, CR1, CR1L, and A2M), and complement initiation pathways: classical (C1QA, C1QB, C1QBP, C1QC, C1R, C1S, C2, C4A, C4B); alternative (CFB, CFD, CFP); and lectin (FCN2, MASP1, MASP2, MBL2). Details of each gene set are provided in Supplementary Table S10, S11. Statistical comparisons between the FSGS and donor groups were performed using Wilcoxon rank-sum tests at the sample level, with Benjamini-Hochberg correction for multiple testing.

### Glomerular pseudotime analysis

To infer disease progression trajectories, pseudotime analysis was performed at the glomerulus level^11,13^ using the quality-controlled dataset prior to cell type reclassification to preserve transcriptomic information and improve the robustness of glomerulus-level pseudotime inference. Gene expression was aggregated across all cells within each glomerulus to generate pseudobulk expression profiles. Dimensionality reduction was performed using Diffusion Map^22^ with k = 10 nearest neighbors, and the first 10 diffusion components were retained for downstream analysis. Pseudotime trajectories were inferred using Slingshot^23^ (version 2.12.0) on the Diffusion Map embedding, with disease group used as cluster labels. The trajectory was constrained to start from the donor group and end at the FSGS group (shrink = 1, extend = “pc1”). Glomeruli were classified into four quartiles (Q1–Q4) based on pseudotime values, representing progression from early (Q1, donor-enriched) to late (Q4, FSGS-enriched) disease states.

### Glomerulus-level differential gene expression analyses

To identify differentially expressed genes between the FSGS and kidney donor glomeruli, pseudobulk expression profiles were generated by aggregating raw transcript counts across the cells within each glomerulus. Differential expression analysis was performed using edgeR^78^ with the quasi-likelihood F-test framework. Lowly expressed genes were filtered using the filterByExpr function, and library sizes were normalized using the trimmed mean of M-values (TMM) method. Dispersions were estimated with robust estimation, and differentially expressed genes were identified using the quasi-likelihood F-test with the contrast FSGS versus kidney donor. Genes with |log FC| > 0.5 and FDR < 0.05 were considered significant. GO enrichment analysis was performed on up– and down-regulated gene sets separately using clusterProfiler (P < 0.05, q-value < 0.05).

### Spatial classification of cells in interstitial region based on glomerular pseudotime

To investigate pseudotime-dependent changes in the interstitial microenvironment, we developed a distance-based spatial classification method. Glomeruli were first classified into quartiles (Q1-Q4) based on their pseudotime values, representing progressive stages of disease. For each cell located in the interstitial region, we calculated the euclidean distance to the centroid of all glomeruli within each quartile group and assigned cells to their nearest glomerular quartile category based on minimum distance.

Within each quartile group, cells in the interstitial region were further stratified into distance-based subcategories using quartiles of the overall distance distribution from all interstitial cells to their nearest glomerulus: Very Close (0-25th percentile of the distance from glomeruli), Close (25-50th percentile), Moderate (50-75th percentile), and Far (75-100th percentile). Prior to distance calculation, tissue regions were manually excluded based on spatial coordinates when they either lacked glomeruli or could be incorrectly assigned to glomeruli from adjacent tissue sections rather than the same section (54,894 cells, 11.1% of total, Extended. Data Fig. 7).

### Spatial visualization of complement pathway scores

For each distance category, complement activation and suppression scores calculated based on SCSE algorism were binarized into “High” and “Low” categories using the median value as the threshold. This categorization was performed separately for each distance category to account for potential distance-dependent variations in score distributions.

To visualize spatial patterns around glomeruli, we generated centralized maps using relative coordinates. For each cell, relative position was calculated as:

Relative X = Cell X coordinate – Nearest glomerulus centroid X

Relative Y = Cell Y coordinate – Nearest glomerulus centroid Y

This transformation centered all glomeruli at the origin (0, 0), enabling direct comparison of spatial patterns across different glomeruli.

### Cell–cell communication analysis

Cell–cell communication analysis was performed with CellChat^27^ (version 2.1.2), using normalized expression data and spatial coordinates to infer spatially constrained ligand– receptor interactions. CellChat objects were constructed separately for FSGS and kidney donor samples, incorporating cell coordinates with a conversion factor of 1 (micrometers) and a cell size of 10 μm. Significant ligand–receptor interactions were inferred using the computeCommunProb function (type = “trimean”, trim = 0.2, distance.use = TRUE, contact.dependent = TRUE, contact.range = 10, interaction.range = 100, scale.distance = 0.2) with the CellChatDB. For comparison between the groups, CellChat objects were merged using the mergeCellChat function. Differentially expressed signaling was identified using a log fold-change threshold of 0.25, a minimum expression fraction of 0.05, and a p-value threshold of 0.05. For periglomerular analysis, cells within glomeruli and the adjacent interstitial region were subset, and glomeruli were further stratified into the pseudotime quartiles (Q1–Q4).

CellChat analyses were performed for each quartile, and the resulting CellChat objects were integrated using mergeCellChat function to assess changes in intercellular communication across disease progression. Spatial visualization of cell–cell communication was performed by integrating CellChat results with Xenium spatial transcriptomics data. Cell and nucleus boundaries were extracted from Xenium parquet files, and communication events were visualized as arrows connecting sender and receiver cells within the interaction range (100 μm).

### Statistical analysis of clinical and spatial transcriptomics data

Statistical analyses for patient characteristics, glomerular percentages, and quantitative immunohistochemical staining were performed using EZR version 1.63 and R version 4.2.2. Kruskal–Wallis test was used for continuous variables and Fisher’s exact test was used for categorical variables. When significant differences were observed, post hoc multiple comparisons were conducted using the Bonferroni correction or Dunn’s test. Two-sided *P* values < 0.05 were considered statistically significant.

For spatial transcriptomics data, differential gene expression was assessed using the Wilcoxon rank-sum test with Benjamini–Hochberg correction for multiple comparisons. Genes were considered differentially expressed when they met the thresholds of min.pct > 0.25, |log FC| > 0.25, and adjusted P < 0.05. Pathway enrichment analysis was performed using ORA based on the list of DEGs identified with thresholds of adjusted P < 0.05 and |log FC| > 0.25. In the glomerulus-level analysis, cell types with fewer than 10 cells across the entire dataset were excluded from correlation analysis. For comparisons of complement scores across pseudotime quartiles within each distance category, the Kruskal-Wallis test was applied at the sample level, followed by Dunn’s post-hoc test with Bonferroni correction when the omnibus test was significant (P < 0.05).

