## Extended Figures for "Multi-Omics Mapping of Human Kidney Reveals Complement-Mediated Cellular Dynamics During Progression of Focal Segmental Glomerulosclerosis"

#### Extended Data Figures

Extended Data Fig. 1. Quality assessment of kidney tissue composition and proteomics data.

Extended Data Fig. 2. Differential protein abundance in FSGS and disease controls.

Extended Data Fig. 3. Eleven proteins elevated in FSGS relative to healthy and disease controls.

Extended Data Fig. 4. Integration, annotation, and refinement of Xenium spatial cell states.

Extended Data Fig. 5. Pseudotime analysis of glomerular cells reveals disease-associated transcriptional trajectories.

Extended Data Fig. 6. Glomerular cell type distribution in FSGS and donors.

Extended Data Fig. 7. Classification of interstitial regions according to glomerular pseudotime and distance.

Extended Data Fig. 8. Cell–cell interaction analyses across disease states and glomerular pseudotime in glomerular and periglomerular regions.

#### Extended Fig. 1

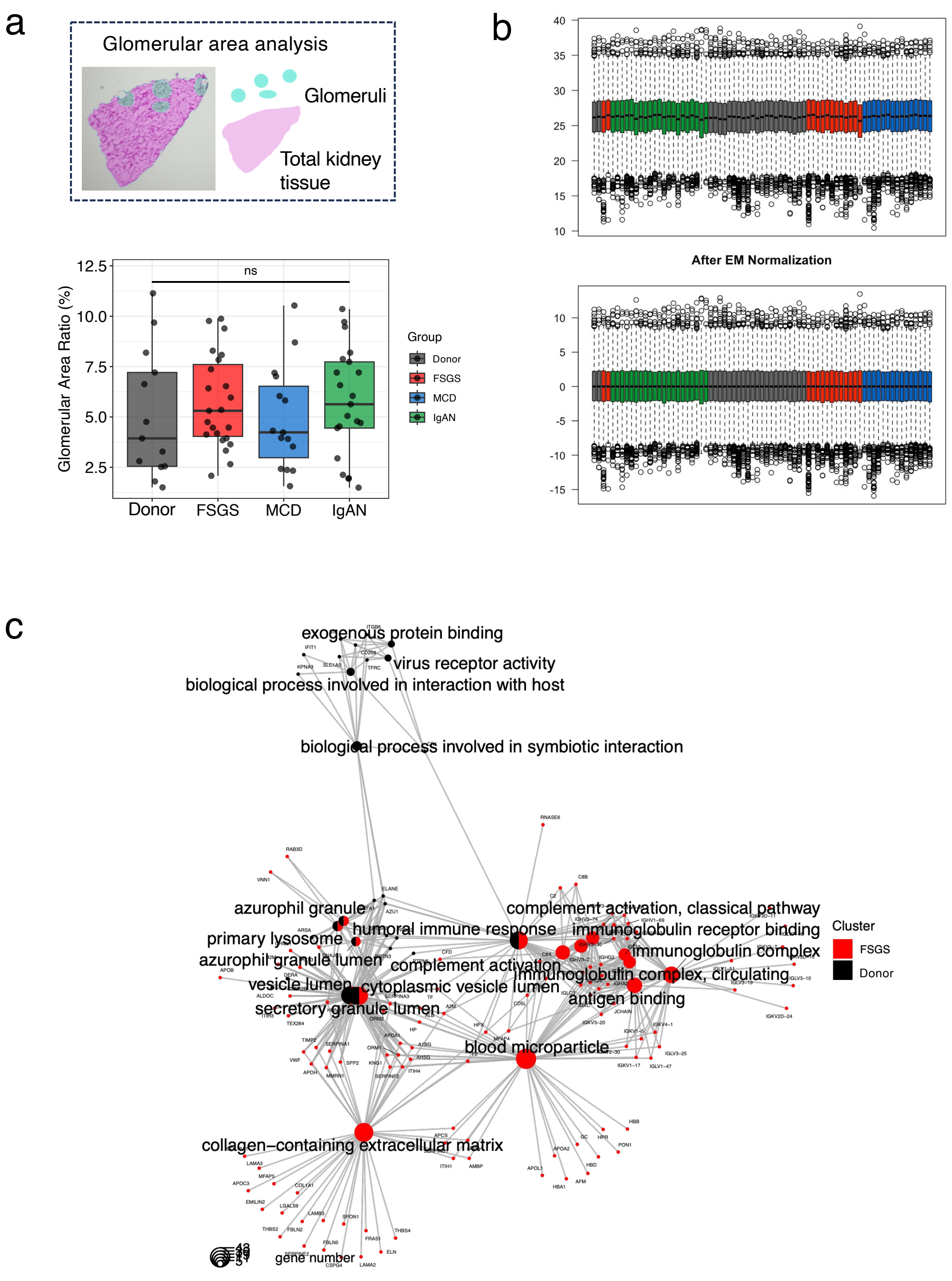


**Extended Data Fig. 1. Quality assessment of kidney tissue composition and proteomics data.**

**a,** Serial sections used for LC–MS/MS are stained with hematoxylin and eosin (HE) to quantify the proportion of glomerular area (light blue) within total kidney tissue area (pink). Glomerular area ratios are compared among the kidney donor, FSGS, MCD, and IgAN groups. Statistical differences are assessed using the Kruskal–Wallis test. ns: not significant. **b,** Quality assessment and distribution of the LC–MS/MS dataset. Data distributions before and after equal median (EM) normalization are shown (top: pre-normalization; bottom: post-normalization). **c,** Over-representation analysis (ORA) of Gene Ontology pathways for the FSGS group compared with the kidney donor group. Pathways and genes upregulated in the FSGS group are shown in red and those in the kidney donor group are shown in black.

### Extended Fig. 2

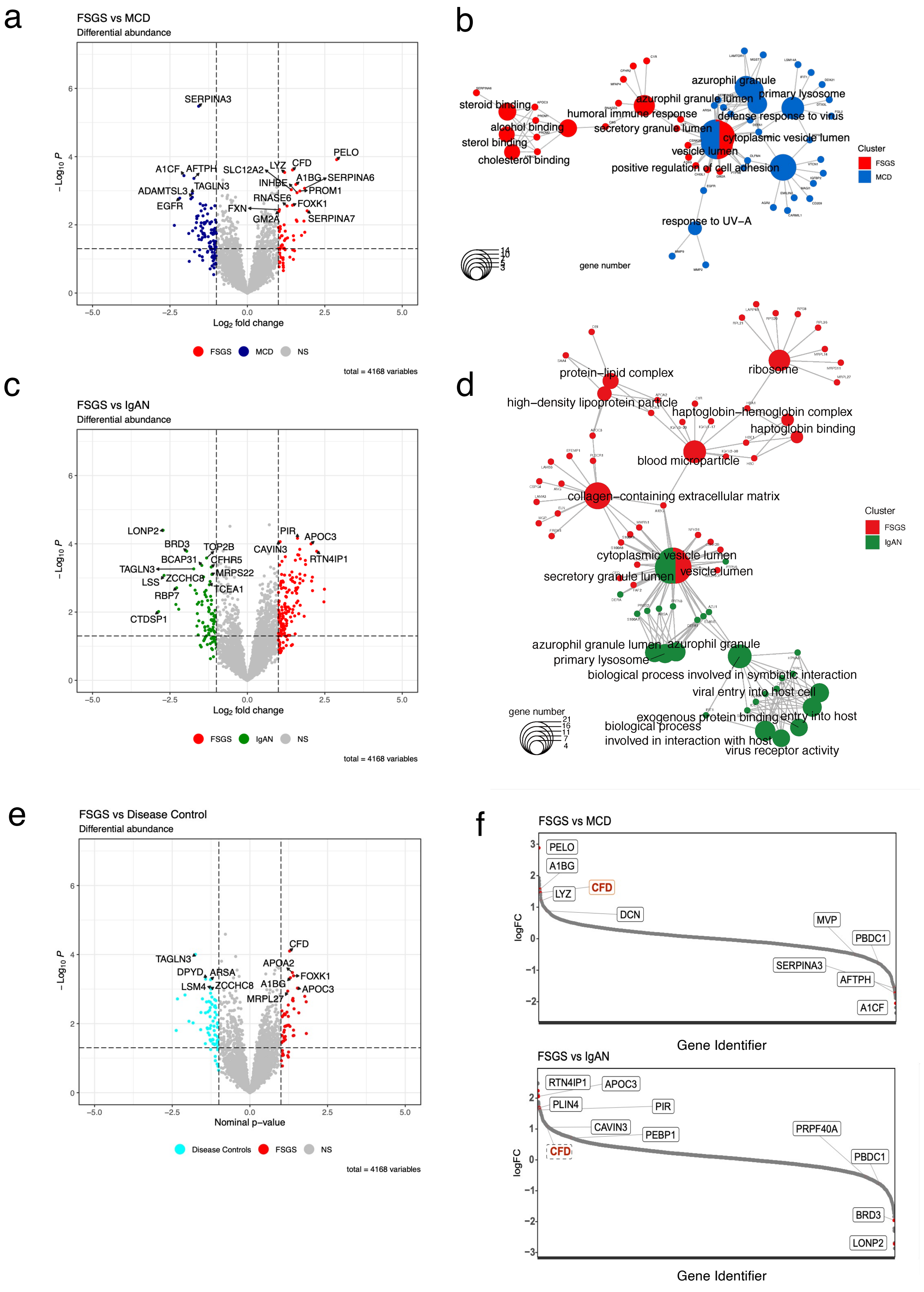


**Extended Data Fig. 2. Differential protein abundance in FSGS and disease controls.**

**a, c, e,** Volcano plots comparing FSGS with each disease group (a, MCD; c, IgAN; e, disease controls [MCD + IgAN]). Significance thresholds are set at $P < 0.05$ and $\log_2$ fold-change > 1.0. **b, d,** Over-representation analysis (ORA) of Gene Ontology pathways comparing the FSGS group with the corresponding disease groups (b, MCD; d, IgAN). Pathways and genes upregulated in FSGS are shown in red, whereas those elevated in MCD and IgAN are shown in blue and green, respectively. **f,** Rank-intensity plots corresponding to the analyses in panels a and c.

#### Extended Fig. 3

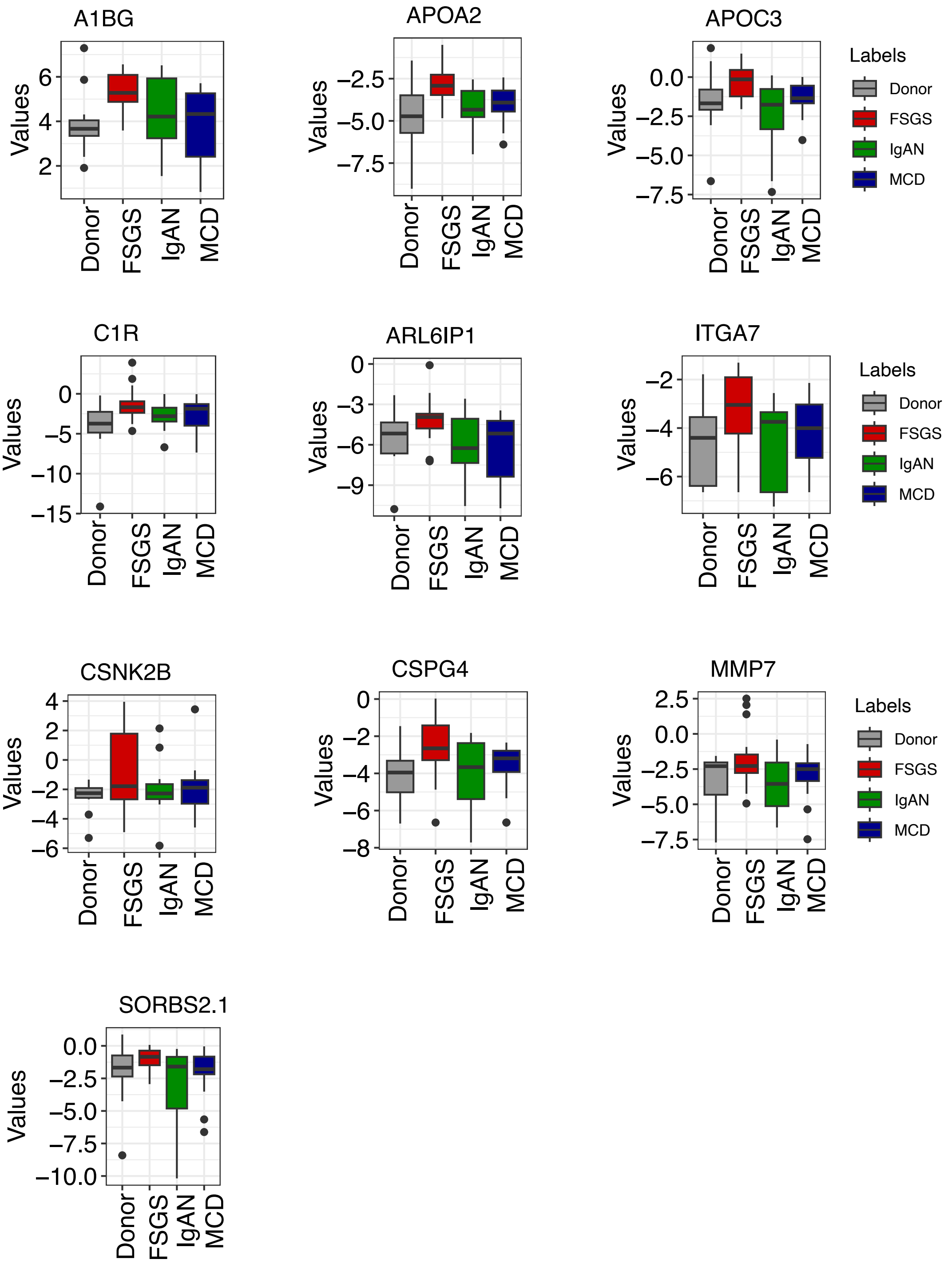


**Extended Data Fig. 3. Eleven proteins elevated in FSGS relative to healthy and disease controls.**

Boxplots of 11 proteins elevated in the FSGS group (excluding CFD) compared with the healthy and disease controls. A1BG: alpha-1B-glycoprotein, APOA2: apolipoprotein A-II, APOC3:

apolipoprotein C-III, C1R: complement C1r subcomponent, ARL6IP1: ADP-ribosylation factor-like protein 6-interacting protein 1, ITGA7: integrin alpha-7, CSNK2B: casein kinase II subunit beta, CSPG4: chondroitin sulfate proteoglycan 4, MMP7: matrilysin, SORBS2: isoform 12 of Sorbin and SH3 domain-containing protein 2

### Extended Fig. 4

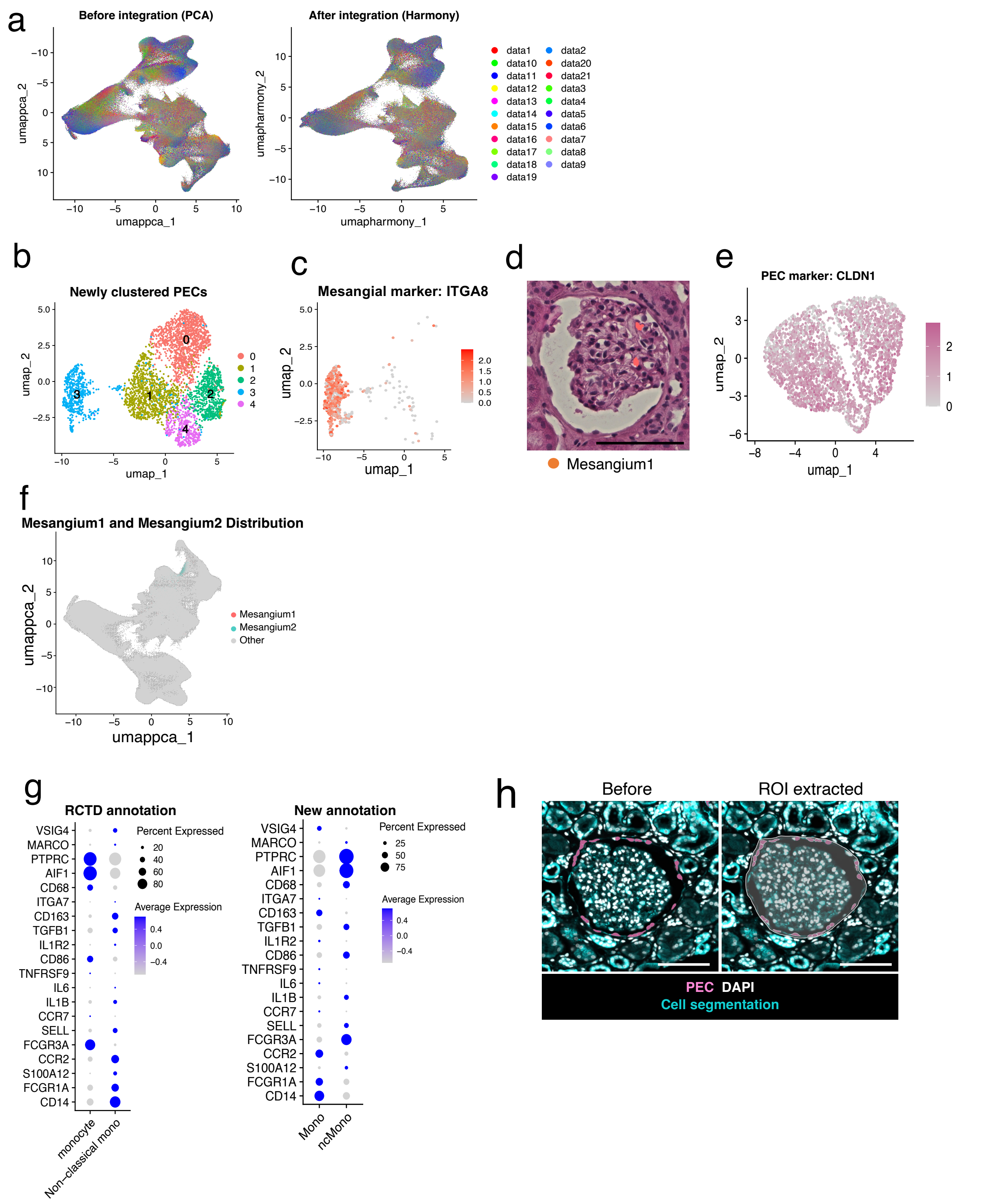


**Extended Data Fig. 4. Integration, annotation, and refinement of Xenium spatial cell states.**

**a**, UMAP plots of all samples before and after dataset integration using PCA (left) and Harmony (right). **b–d,** Subclustering of PECs and identification of Mesangial cells. **b,** UMAP plot after removal of extraglomerular PECs, highlighting refined PEC-associated clusters. **c,** Cluster 3 shows high expression of the mesangial marker ITGA8 and is annotated as Mesangium 1. **d,** Representative HE-stained image showing the location of Mesangium 1 cells within a mesangial region (scale bar, 100 μm). **e,** UMAP plot after exclusion of Mesangium 1, demonstrating that the remaining PEC population robustly expresses the PEC marker CLDN1. **f,** UMAP projection showing the distribution of Mesangium 1 and Mesangium 2 clusters across all cells. **g,** Non-classical monocytes lacking FCGR3A (CD16) are distinguished from classical monocytes to ensure accurate subpopulation identification. **h,** Representative images after extraction of a glomerular region of interest (ROI). Left, original image before extraction. Right, extracted glomerular ROI delineated by a boundary.

PEC, parietal epithelial cell. Scale bars, 100 μm.

#### Extended Fig. 5

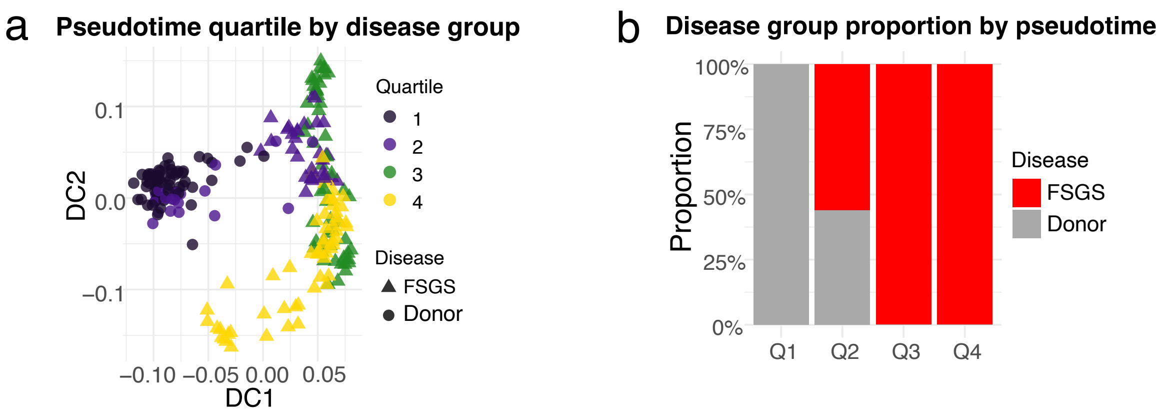


**Extended Data Fig. 5. Pseudotime analysis of glomerular cells reveals disease-associated transcriptional trajectories.**

**a,** Diffusion map of glomeruli coloured by disease status and pseudotime. Glomeruli from the FSGS group are shown as triangles and those from kidney donors as circles. Pseudotime is indicated by colour. **b,** Distribution of glomeruli from each group across pseudotime quartiles (Q1–Q4).

#### Extended Fig. 6

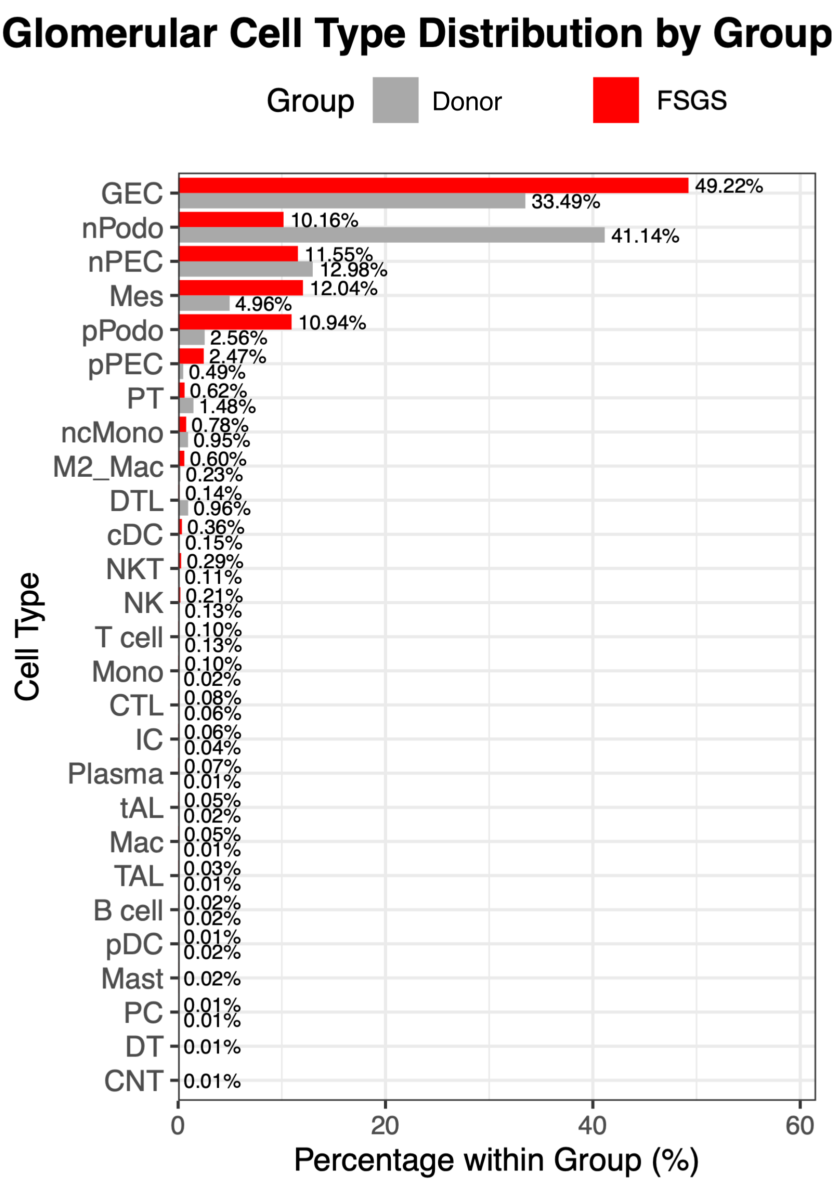


**Extended Data Fig. 6. Glomerular cell type distribution in FSGS and donors.**

Comparison of the percentages of cell types detected in the glomeruli between the FSGS and kidney donor groups.

#### Extended Fig. 7

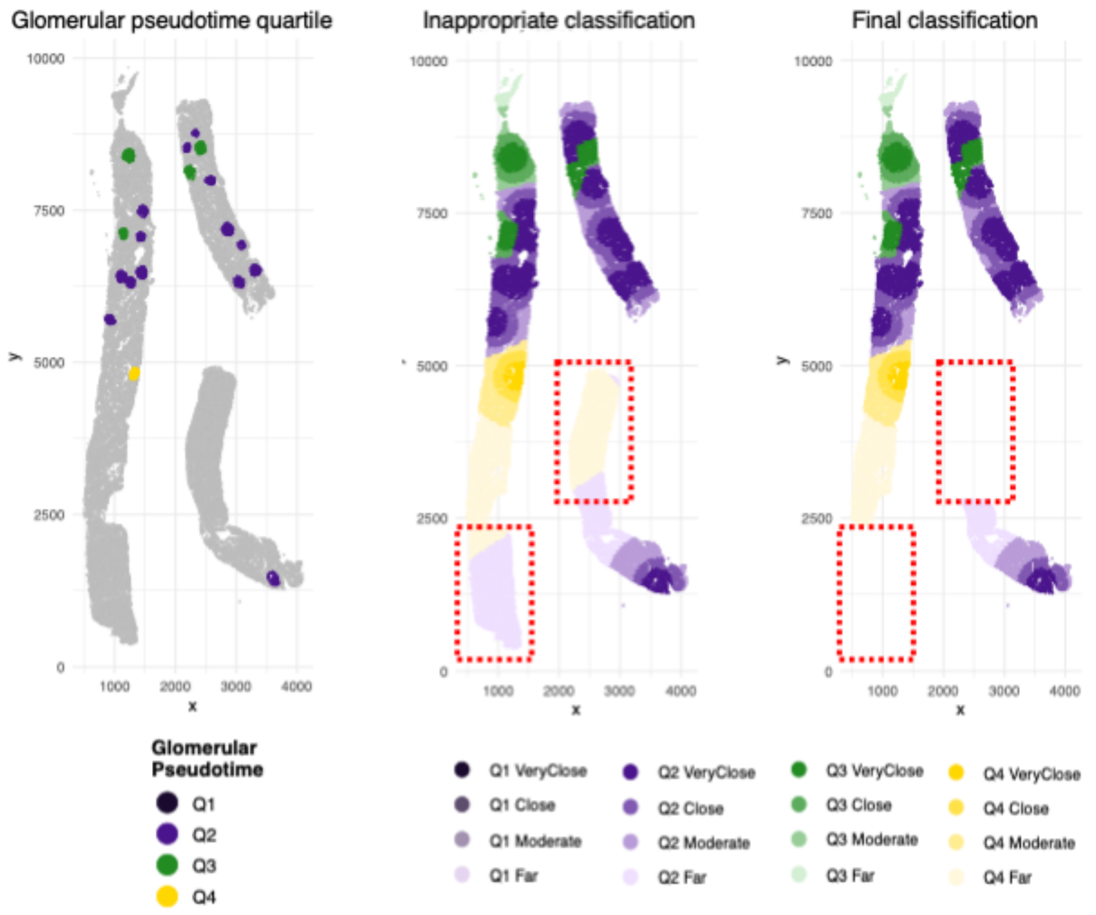


**Extended Data Fig. 7. Classification of interstitial regions according to glomerular pseudotime and distance**

Representative spatial maps of interstitial regions relative to glomerular pseudotime and distance. Left, glomeruli coloured by pseudotime quartiles. Middle, cells in the interstitial region assigned to the nearest glomerular quartile based on Euclidean distance. Right, cells incorrectly assigned to quartiles are excluded to refine the mapping. Red dotted squares indicate inappropriate mappings.

### Extended Fig. 8

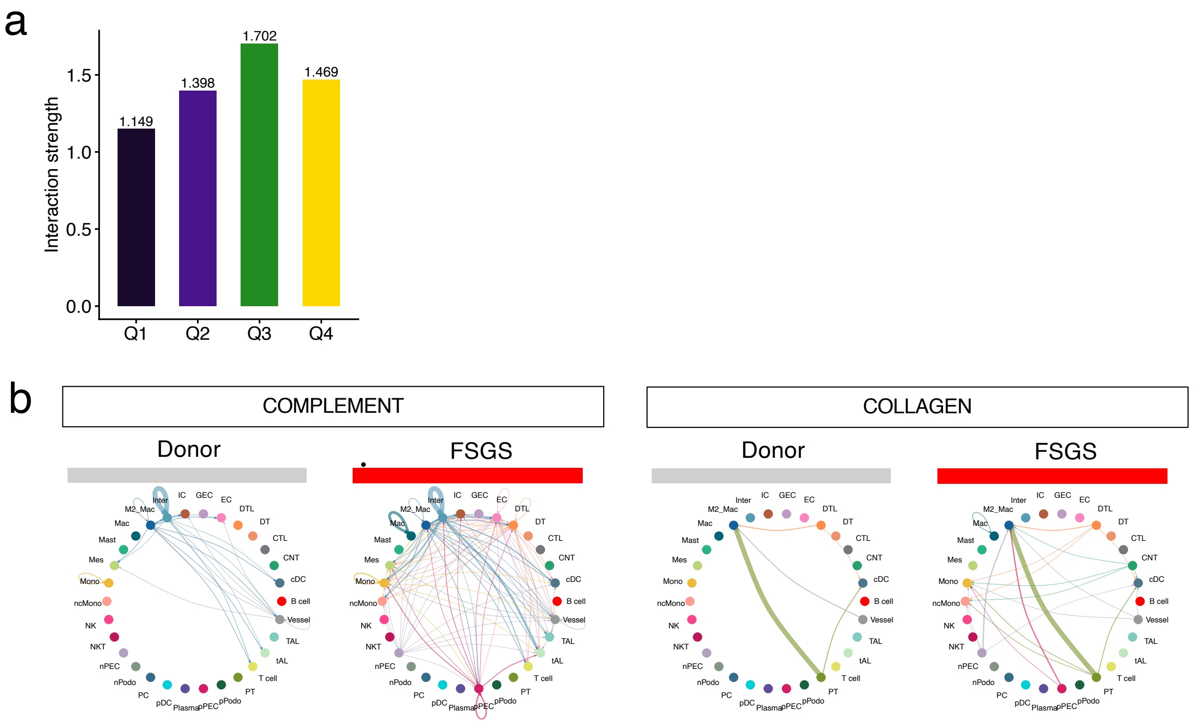


**Extended Data Fig. 8. Cell–cell interaction analyses across disease states and glomerular pseudotime in glomerular and periglomerular regions.**

**a,** Bar plots showing the strength of cell-cell interactions in each glomerular pseudotime quartile. **b,** Circle plots showing intercellular interactions of complement (left) and collagen (right) signals. Cell type abbreviations: DTL, thin descending limb; IC, intercalated cell; PT, proximal tubule; Inter, interstitial cell; CNT, connecting tubule; PC, principal cell; TAL, thick ascending limb; DT, distal tubule; cDC, conventional dendritic cell; tAL, thin ascending limb; nPEC, normal parietal epithelial cell; pPEC, pathogenic parietal epithelial cell; nPodo, normal podocyte; pPodo, pathogenic podocyte; GEC, glomerular endothelial cell; CTL, cytotoxic T cell; ncMono, non-classical monocyte; M2_Mac, M2 macrophage; EC, extraglomerular endothelial cell; Mes, mesangium; Mono, monocyte; Mac, macrophage; pDC, plasmacytoid dendritic cell
