## Supplementary information for "Multi-Omics Mapping of Human Kidney Reveals Complement-Mediated Cellular Dynamics During Progression of Focal Segmental Glomerulosclerosis"

### Methods

#### LC-MS/MS data acquisition

Full MS scans were acquired at a resolution of 70,000 (automatic gain control (AGC) target of 3e6, max injection time of 100 ms, m/z range 350-1500). MS/MS scans were conducted at a resolution of 17,500 (AGC target of 1e5, max injection time of 50 ms, m/z range 200-2000), selecting the top 10 precursor ions with an isolation window of 2.0 m/z. Dynamic exclusion was set to auto, with a minimum AGC target of 1.00e3 and an intensity threshold of 2.0e4. Precursor ions with charge states of 1, 5, more than 5, or unassigned were excluded. Isotope exclusion was enabled. The spectra were imported into Proteome Discoverer 2.5 (PD 2.5; Thermo Fisher Scientific) as follows: the four SDB-SCX fractions per kidney biopsy sample were imported as 'fractions' thereby combining them into a single query file; for a total of 73 queries that were analyzed against the Uniprot fasta database (downloaded March 16, 2023; n = 42,326 entries) using the Sequest HT algorithm. Trypsin was set as the digestion enzyme, allowing up to 2 missed cleavages and a minimum peptide length of 6 amino acids. Oxidation (+15.995 Da) of methionine; acetylation (+42.011 Da) of the N-terminus; the loss of N-terminal methionine (-131.040 Da); and combining loss of methionine and addition of acetylation at N-terminal methionine (-89.030 Da), were set as dynamic modifications. Spectral mass tolerances were 10 ppm for the precursor and 20 mmu for the fragments.

**Supplementary table S1. The patients' background at kidney biopsy in the proteomics cohort.**

| Total<br>n = 73 | FSGS<br>n = 23 | Donor<br>n = 14 | MCD<br>n = 15 | IgAN<br>n = 21 | P<br>value |
|---|---|---|---|---|---|
| Sex (%) | [a)] | | [d)] | [e)] | 0.028 |
| Female | 8 (34.8) | 10 (76.9) | 4 (26.7) | 7 (33.3) | |
| Male | 15 (65.2) | 3 (23.1) | 11 (73.3) | 14 (66.7) | |
| Age | 63.00 | 66.00 | 70.00 | 64.00 | 0.549 |
| | [44.50, 71.00] | [45.00, 67.00] | [52.50, 72.50] | [39.00, 69.00] | |
| BMI | 24.60 | 22.54 | 25.09 | 23.36 | 0.64 |
| ($kg/m^2$) | [21.53, 27.17] | [21.25, 27.12] | [22.84, 28.69] | [21.70, 27.18] | |
| Mean BP | 100.00 | 90.33 | 90.00 | 98.67 [e)] | 0.012 |
| (mmHg) | [93.33, 105.17] | [81.50, 95.67] | [83.83, 99.17] | [94.00, 107.00] | |
| TP | 6.50 [b)] | 7.00 | 4.70 [d)] | 6.90 [f)] | <0.001 |
| (g/dl) | [5.50, 7.25] | [6.75, 7.15] | [4.30, 5.00] | [6.50, 7.20] | |
| Alb | 3.80 [a)] [b)] | 4.30 | 1.70 [d)] | 4.10 [f)] | <0.001 |
| (g/dl) | [2.90, 4.15] | [4.10, 4.55] | [1.45, 2.05] | [3.50, 4.10] | |
| T.chol | 231.00 [b)] | 195.00 | 356.00 [d)] | 190.00 [f)] | <0.001 |
| (g/dl) | [204.25, 280.50] | [164.50, 233.50] | [296.00, 469.00] | [171.50, 229.00] | |
| LDL | 146.70 [b)] | 95.60 | 214.00 [d)] | 110.70 [f)] | <0.001 |
| (mg/dl) | [92.40, 183.05] | [90.00, 133.10] | [184.80, 363.20] | [80.15, 126.05] | |
| CRP | 0.05 | 0.08 | 0.08 | 0.07 | 0.947 |
| (mg/dl) | [0.02, 0.22] | [0.03, 0.18] | [0.04, 0.10] | [0.02, 0.10] | |
| Cr | 1.16 [a)] | 0.62 | 1.02 [d)] | 1.08 [e)] | <0.001 |
| (mg/dl) | [1.05, 1.42] | [0.58, 0.67] | [0.84, 1.73] | [0.88, 1.52] | |
| eGFR | 44.94 [a)] | 75.66 | 53.98 [d)] | 50.25 [e)] | 0.001 |
| ($ml/min/1.73m^2$) | [37.35, 55.19] | [70.69, 85.99] | [31.37, 80.86] | [40.70, 65.92] | |
| UP/Ucr | 2.50 [a)] [b)] | 0.06 | 10.15 [d)] | 0.98 [e)] [f)] | <0.001 |
| (g/gCr) | [1.48, 7.92] | [0.06, 0.07] | [8.15, 14.28] | [0.50, 2.72] | |

Data are expressed as Median [interquartile range] or numbers (%).

Statistic test: Kruskal-Wallis test for numeric data, and Fisher's exact test for categorical variables.

P values indicate the results of the Kruskal-Wallis test or Fisher's exact test performed on the four groups. P values less than 0.05 were considered statistically significant. If there are any significant differences, multiple tests were performed by Bonferroni method. When any significant differences are detected, following a), b), c), d), e) and f) are shown.

a) FSGS – Kidney Donor, b) FSGS – MCD, c) FSGS – IgAN, d) Kidney Donor – MCD, e) Kidney Donor – IgAN, f) MCD – IgAN

FSGS: focal segmental glomerulosclerosis, MCD: minimal change disease, IgAN: IgA nephropathy, BMI: body mass index, Mean BP: mean blood pressure, TP: serum total protein, Alb: serum albumin, T.chol: serum total cholesterol, LDL: serum low density lipoprotein cholesterol, CRP: serum c reactive protein, Cr: serum creatinine, eGFR: estimated glomerular filtration rate, UP/Ucr: urinary protein/ urinary creatinine ratio

**Supplementary table S2. The patients' background at kidney biopsy in the spatial transcriptomics cohort.**

| **Total**<br>**n = 21** | **FSGS**<br>**n = 15** | **Donor**<br>**n = 6** | **P**<br>**value** |
|---|---|---|---|
| Sex (%) | | | 0.028 |
| Female | 6 ( 40.0) | 4 ( 66.7) | |
| Male | 9 ( 60.0) | 2 ( 33.3) | |
| Age | 60.00<br>[52.00, 74.00] | 44.50<br>[38.00, 62.25] | 0.129 |
| BMI<br>(kg/m$^2$) | 23.60<br>[22.89, 27.72] | 21.86<br>[19.92, 25.43] | 0.096 |
| Mean BP<br>(mmHg) | 103.33<br>[89.42, 108.83] | 94.33<br>[89.08, 97.33] | 0.386 |
| TP<br>(g/dl) | 5.40<br>[4.85, 6.80] | 7.30<br>[7.12, 7.47] | 0.001 |
| Alb<br>(g/dl) | 2.50<br>[2.05, 3.75] | 4.40<br>[4.18, 4.40] | 0.002 |
| T.chol<br>(g/dl) | 283.00<br>[236.25, 325.00] | 169.00<br>[166.50, 204.50] | 0.015 |
| LDL<br>(mg/dl) | 180.65<br>[142.50, 206.00] | 93.50<br>[81.50, 119.00] | 0.017 |
| CRP<br>(mg/dl) | 0.05<br>[0.01, 0.18] | 0.03<br>[0.02, 0.07] | 0.557 |
| Cr<br>(mg/dl) | 1.05<br>[0.80, 1.54] | 0.64<br>[0.63, 0.84] | 0.087 |
| eGFR<br>(ml/min/1.73m$^2$) | 48.72<br>[31.22, 74.41] | 74.50<br>[70.20, 80.83] | 0.073 |
| UP/Ucr<br>(g/gCr) | 6.41<br>[3.08, 10.86] | 0.06<br>[0.05, 0.08] | <0.001 |

Data are expressed as Median [interquartile range] or numbers (%).

Statistic test: Mann-Whitney U test for numeric data, and Fisher's exact test for categorical variables

P-values less than 0.05 were considered statistically significant.

FSGS: focal segmental glomerulosclerosis, BMI: body mass index, Mean BP: mean blood pressure, TP: serum total protein, Alb: serum albumin, T.chol: serum total cholesterol, LDL: serum low density lipoprotein cholesterol, CRP: serum c reactive protein, Cr: serum creatinine, eGFR: estimated glomerular filtration rate, UP/Ucr: urinary protein/ urinary creatinine ratio

**Supplementary table S3. Classification, treatment, and electron microscopic findings of FSGS groups used in the proteomics and spatial transcriptomics cohorts.**

| | **Proteomics cohort n = 23** | | **Spatial cohort n = 15** | |
|---|---|---|---|---|
| | Number of patients | % | Number of patients | % |
| Primary FSGS | 9 | 39.1 | 10 | 66.7 |
| Secondary FSGS | 8 | 34.8 | 0 | 0.0 |
| Undetermined cause | 6 | 26.1 | 5 | 33.3 |
| Steroid therapy | 11 | 47.8 | 9 | 60.0 |
| Immunosuppressants therapy | 4 | 17.4 | 2 | 13.3 |
| Foot process effacement | | | | |
| Diffuse | 5 | 21.7 | 8 | 53.3 |
| Focal | 4 | 17.4 | 3 | 20.0 |
| Negative | 0 | 0.0 | 1 | 6.7 |
| Not available | 14 | 60.9 | 3 | 20.0 |

Data are expressed as Number of patients or percentages (%).

FSGS: focal segmental glomerulosclerosis

**Supplementary table S4. Evaluation of pathological differences in the kidney biopsy samples among the four groups in proteomics cohort.**

| **Total n = 73** | **FSGS n = 23** | **Donor n = 14** | **MCD n = 15** | **IgAN n = 21** | **P value** |
|---|---|---|---|---|---|
| Global sclerosis (%) | 31.25 [a) b)] [13.10, 49.15] | 6.70 [0.00, 10.44] | 11.11 [6.38, 23.08] | 17.39 [e)] [11.11, 26.32] | 0.001 |
| Segmental sclerosis (%) | 9.09 [a) b) c)] [4.88, 17.42] | 0.00 [0.00, 0.00] | 0.00 [0.00, 0.00] | 0.00 [0.00, 4.35] | <0.001 |
| Interstitium cell infiltration (%) | [a)] | | | [f) e)] | 0.047 |
| None | 13 (56.5) | 14 (100.0) | 12 (80.0) | 8 (38.1) | |
| Mild | 7 (30.4) | 0 (0.0) | 3 (20.0) | 10 (47.6) | |
| Moderate | 2 (8.7) | 0 (0.0) | 0 (0.0) | 2 (9.5) | |
| Severe | 1 (4.3) | 0 (0.0) | 0 (0.0) | 1 (4.8) | |
| Interstitium fibrosis (%) | [a) b)] | | | [e) f)] | <0.001 |
| None | 5 (21.7) | 14 (100.0) | 11 (73.3) | 4 (19.0) | |
| Mild | 13 (56.5) | 0 (0.0) | 3 (20.0) | 10 (47.6) | |
| Moderate | 4 (17.4) | 0 (0.0) | 1 (6.7) | 7 (33.3) | |
| Severe | 1 (4.3) | 0 (0.0) | 0 (0.0) | 0 (0.0) | |
| IF C3 | [c)] | | | [e) f)] | <0.001 |
| (-) | 22 (95.7) | 12 (85.7) | 15 (100.0) | 5 (23.8) | |
| (+) | 0 (0.0) | 2 (14.3) | 0 (0.0) | 13 (61.9) | |
| (++) | 1 (4.3) | 0 (0.0) | 0 (0.0) | 3 (14.3) | |
| IF Fib | [c)] | | | [e) f)] | <0.001 |
| (-) | 21 (91.3) | 14 (100.0) | 15 (100.0) | 10 (47.6) | |
| (+) | 2 (8.7) | 0 (0.0) | 0 (0.0) | 9 (42.9) | |
| (++) | 0 (0.0) | 0 (0.0) | 0 (0.0) | 2 (9.5) | |
| IF IgA | [c)] | | | [e) f)] | <0.001 |
| (-) | 20 (87.0) | 10 (71.4) | 14 (93.3) | 0 (0.0) | |
| (+) | 3 (13.0) | 4 (28.6) | 1 (6.7) | 8 (38.1) | |
| (++) | 0 (0.0) | 0 (0.0) | 0 (0.0) | 9 (42.9) | |
| (+++) | 0 (0.0) | 0 (0.0) | 0 (0.0) | 4 (19.0) | |
| IF IgG | | | | | 0.422 |
| (-) | 22 (95.7) | 14 (100) | 15 100.0() | 19 (90.5) | |

| | | | | | |
|---|---|---|---|---|---|
| (+) | 1 (4.3) | 0 (0.0) | 0 (0.0) | 2 (9.5) | |
| IF IgM | | | | | 0.311 |
| (-) | 18 (78.3) | 14 (100.0) | 13 (86.7) | 17 (81.0) | |
| (+) | 5 (21.7) | 0 (0.0) | 2 (13.3) | 4 (19.0) | |
| IF kappa chain | [c)] | | | [e) f)] | <0.001 |
| (-) | 21 (91.3) | 14 (100.0) | 15 (100.0) | 3 (14.3) | |
| (+) | 2 (8.7) | 0 (0.0) | 0 (0.0) | 15 (71.4) | |
| (++) | 0 (0.0) | 0 (0.0) | 0 (0.0) | 3 (14.3) | |
| IF lamda chain | [c)] | | | [e) f)] | <0.001 |
| (-) | 21 (91.3) | 12 (85.7) | 15 (100.0) | 3 (14.3) | |
| (+) | 2 (8.7) | 2 (14.3) | 0 (0.0) | 14 (66.7) | |
| (++) | 0 (0.0) | 0 (0.0) | 0 (0.0) | 4 (19.0) | |

Data are expressed as Median [interquartile range] or numbers (%).

Statistic test: Kruskal-Wallis test for numeric data, and Fisher‘s exact test for categorical variables.

P-values indicate the results of the Kruskal-Wallis test or Fisher‘s exact test performed on the four groups. P-values less than 0.05 were considered statistically significant. If there are any significant differences, multiple tests were performed by Bonferroni method. When any significant differences are detected, following a), b), c), d), e) and f) are shown.

a) FSGS - Kidney Donor, b) FSGS – MCD, c) FSGS – IgAN, d) Kidney Donor – MCD, e) Kidney Donor – IgAN, f) MCD – IgAN

Interstitial cell infiltration, interstitial fibrosis are defined as follows: None: <10% of cortical area, Mild: 10–24%, Moderate: 25–49%, Severe: >50%.

FSGS: focal segmental glomerulosclerosis, MCD: minimal change disease, IgAN: IgA nephropathy

**Supplementary table S5. Evaluation of pathological differences in the kidney biopsy samples among the two groups in spatial transcriptomics cohort.**

| Total<br>n = 21 | FSGS<br>n = 15 | Donor<br>n = 6 | P<br>value |
|---|---|---|---|
| Global sclerosis (%) | 15.79<br>[7.99, 45.69] | 2.38<br>[0.00, 20.93] | 0.086 |
| Segmental sclerosis (%) | 9.09<br>[3.24, 11.81] | 0.00<br>[0.00, 0.00] | 0.004 |
| Interstitium cell infiltration (%) | | | 0.156 |
| None | 8 (53.3) | 6 (100.0) | |
| Mild | 6 (40.0) | 0 (0.0) | |
| Moderate | 1 (6.7) | 0 (0.0) | |
| Interstitium fibrosis (%) | | | 0.29 |
| None | 5 (33.3) | 5 (83.3) | |
| Mild | 6 (40.0) | 1 (16.7) | |
| Moderate | 3 (20.0) | 0 (0.0) | |
| Severe | 1 (6.7) | 0 ( 0.0) | |
| IF C3 | [c)] | | 1 |
| (-) | 14 (93.3) | 6 (100.0) | |
| (+) | 1 (6.7) | 0 (0.0) | |
| IF Fib | | | N/A |
| (-) | 15 (100.0) | 6 (100.0) | |
| IF IgA | | | 0.167 |
| (-) | 12 (80.0) | 5 (83.3) | 0.167 |
| (±) | 3 (20.0) | 0 (0.0) | |
| (+) | 0 (0.0) | 1 (16.7) | |
| IF IgG | | | N/A |
| (-) | 15 (100.0) | 6 (100.0) | |
| IF IgM | | | 0.43 |
| (-) | 13 (86.7) | 4 (66.7) | |
| (±) | 0 (0.0) | 1 (16.7) | |
| (+) | 2 (13.3) | 1 (16.7) | |
| IF kappa chain | | | N/A |
| (-) | 15 (100.0) | 6 (100.0) | |

| | | | |
|---|---|---|---|
| IF lamda chain | | | N/A |
| (-) | 15 (100.0) | 6 (100.0) | |

Data are expressed as Median [interquartile range] or numbers (%).

Statistic test: Kruskal-Wallis test for numeric data, and Fisher's exact test for categorical variables.

P-values indicate the results of the Kruskal-Wallis test or Fisher's exact test performed on the four groups. P-values less than 0.05 were considered statistically significant. N/A indicates not applicable.

If there are any significant differences, multiple tests were performed by Bonferroni method. When any significant differences are detected, following a), b), c), d), e) and f) are shown.

a) FSGS - Kidney Donor, b) FSGS – MCD, c) FSGS – IgAN, d) Kidney Donor – MCD, e) Kidney Donor – IgAN, f) MCD – IgAN

Interstitial cell infiltration, interstitial fibrosis are defined as follows: None: <10% of cortical area, Mild: 10–24%, Moderate: 25–49%, Severe: >50%.

FSGS: focal segmental glomerulosclerosis, MCD: minimal change disease, IgAN: IgA nephropathy
